# Nanobody-directed targeting of optogenetic tools to study signaling in the primary cilium

**DOI:** 10.1101/2020.02.04.933440

**Authors:** Jan N. Hansen, Fabian Kaiser, Christina Klausen, Birthe Stüven, Raymond Chong, Wolfgang Bönigk, David U. Mick, Andreas Möglich, Nathalie Jurisch-Yaksi, Florian I. Schmidt, Dagmar Wachten

## Abstract

Compartmentalization of cellular signaling forms the molecular basis of cellular behavior. The primary cilium constitutes a subcellular compartment that orchestrates signal transduction independent from the cell body. Ciliary dysfunction causes severe diseases, termed ciliopathies. Analyzing ciliary signaling and function has been challenging due to the lack of tools to temporarily manipulate and analyze ciliary signaling. Here, we describe a nanobody-based targeting approach for optogenetic tools that is applicable *in vitro* and *in vivo* and allows to specifically analyze ciliary signaling and function. Thereby, we overcome the loss of protein function observed after direct fusion to a ciliary targeting sequence. We functionally localized modifiers of cAMP signaling, i.e. the photo-activated adenylate cyclase bPAC and the light-activated phosphodiesterase LAPD, as well as the cAMP biosensor mlCNBD-FRET to the cilium. Using this approach, we studied the contribution of spatial cAMP signaling in controlling cilia length. Combining optogenetics with nanobody-based targeting will pave the way to the molecular understanding of ciliary function in health and disease.

## Introduction

Primary cilia are membrane-encased protrusions that extend from the surface of almost all vertebrate cells. Primary cilia function as antennae that translate sensory information into a cellular response. The sensory function is governed by a subset of receptors and downstream signaling components that are specifically targeted to the cilium. This allows to orchestrate rapid signal transduction in a minuscule reaction volume, independent of the cell body. A central component of ciliary signaling is the second messenger 3’, 5’-cyclic adenosine monophosphate (cAMP) [1]. The prime example is chemosensation in highly specialized olfactory cilia: odorant-induced activation of G protein-coupled receptors (GPCRs) stimulates the synthesis of cAMP by the transmembrane adenylate cyclase 3 (AC3). The ensuing increase in ciliary cAMP levels activates cyclic nucleotide-gated ion channels (CNG), resulting in a depolarization that spreads from the cilium to the synapse [2]. In recent years, it emerged that cAMP also controls signaling in primary cilia. AC3 is highly enriched in primary cilia and widely used as a ciliary marker [3, 4]. Loss-of-function mutations in the *ADCY3* gene, encoding for AC3, or loss of *ADCY3* expression cause monogenic severe obesity and increase the risk for type 2 diabetes [5–10]. This has been attributed to the loss of AC3 function in neuronal primary cilia [9, 11]. Furthermore, the most prominent primary cilia signaling pathway, the Sonic hedgehog (Shh) pathway, utilizes cAMP as a second messenger in the cilium to transduce stimulation by Shh into a change in gene expression [12, 13]. Finally, the dynamic modulation of primary cilia length seems to be controlled by cAMP [14–16]. However, as of yet, it has been impossible to manipulate cAMP dynamics in primary cilia independently from the cell body. Hence, the molecular details and dynamics of cAMP-signaling pathways in primary cilia remain largely unknown.

Optogenetics might be the key to overcome this issue, not least because it has proven to be a powerful method to manipulate and monitor cAMP dynamics in mouse sperm flagella, a specialized motile cilium [17–19]. The photo-activated adenylate cyclase bPAC [20] has been employed to increase flagellar cAMP levels by blue light [19], and the FRET-based cAMP biosensor mlCNBD-FRET has been used to monitor cAMP dynamics in sperm flagella [18]. This cAMP tool kit has been complemented with the red light-activated phosphodiesterase LAPD that allows to decrease cAMP levels in a light-dependent manner [21, 22]. For primary cilia, the challenge is to specifically target these tools to the cilium to investigate cAMP signaling independent from the cell body. Free diffusion of proteins into the primary cilium is limited by the transition zone (TZ) at the base of the cilium [23]. Protein transport into and out of the cilium relies on the intraflagellar transport (IFT) machinery in combination with the BBSome, a multi-protein complex at the ciliary base [24–27]. The combined action of IFT, BBSome, and TZ shape the unique ciliary protein composition [28]. To localize a given optogenetic tool to the primary cilium, the ciliary transport machinery needs to be hijacked. Common strategies involve direct fusion to the C terminus of either a full-length GPCR, e.g. the somatostatin receptor 3 (Sstr3) [24], or a truncated ciliary protein, e.g. the first 201 amino acids (aa) of the ciliary mouse Nphp3 (nephrocystin 3) protein [29, 30]. Fusion of Sstr3 to the N terminus of bPAC or mlCNBD-FRET has already been applied [18, 31]. However, using a full-length GPCR as a fusion partner for ciliary localization might increase the abundance of this specific receptor in the cilium and, thereby, distort the ciliary set of signal receptors. This is particularly disadvantageous when analyzing ciliary cAMP signaling because fusion to GPCRs, which couple to cAMP signaling, might alter basal cAMP levels due to constitutive activity, as has been shown for the commonly used 5-HT_6_ receptor [32]. In contrast, the truncated version of mNphp3 is sufficient to convey ciliary targeting [29], but lacks any other known functionality, whereby its fusion for ciliary targeting represents by large a less invasive targeting approach than using a complete GPCR. Thus, to target optogenetic tools to the primary cilium, we fused aa 1-201 of mNphp3 to their N terminus. To our surprise, this approach largely failed for LAPD and mlCNBD-FRET, because N-terminal fusion disrupts their optogenetic and biosensoric function, respectively. Therefore, we developed a new approach to hijack the ciliary transport machinery to target intracellular nanobodies. The nanobody-based approach allows to specifically target proteins to the primary cilium without directly fusing them to a ciliary protein. Instead, the nanobody is cilia-targeted and tows the proteins of interest to the cilium by binding to a tag contained in the protein of interest.

We show that the cilia-targeted nanobodies bind to eGFP or mCherry-containing proteins of interest in the cytosol, such as bPAC, LAPD, or mlCNBD-FRET. As a complex, the proteins then translocate into the primary cilium. Nanobody binding does not substantially impair protein function, i.e. light-dependent activation of bPAC and LAPD, or cAMP-induced conformational changes of mlCNBD-FRET. We demonstrate the validity of this approach both *in vitro* and *in vivo* in mammalian cells and zebrafish, respectively. Moreover, using nanobody-based ciliary targeting of bPAC, we study the role of ciliary versus cytosolic (cell body) cAMP signaling in controlling cilia length. Our approach principally extends to the ciliary targeting of any protein of interest, which is recognized by a nanobody. Thereby, the huge variety of genetically-encoded tools to manipulate cellular signaling, e.g. membrane potential, protein-protein interactions, or enzymatic activities, and to monitor cellular signaling, e.g. dynamics of Ca^2+^, pH, or the membrane potential, could be exclusively targeted to the primary cilium using our nanobody-based targeting approach. This strategy opens up new avenues for cilia biology and allows to address long-standing questions in cell biology.

## Results

### N-terminal fusion of optogenetic tools interferes with photoactivation

To utilize the optogenetic tools bPAC- or LAPD-mCherry in a cilium-specific manner, we first tested whether N-terminal fusion of mNphp3(201) is sufficient for targeting to the primary cilium. Indeed, fusions of both bPAC and LAPD predominantly localized to the primary cilium (Fig. 1A, B). To test whether protein fusion interferes with the light-dependent activation of LAPD or bPAC, we measured LAPD or bPAC activity using Ca^2+^ imaging. To this end, we used HEK293 cells stably expressing a cyclic nucleotide-gated (CNG) ion channel CNGA2-TM (HEK-TM) that conducts Ca^2+^ upon cAMP binding [33]. HEK293 cells were not ciliated to directly compare the non-fused and the fused optogenetic tool. Activation of bPAC with a 465 nm light pulse increases intracellular cAMP levels, leading to a Ca^2+^ influx, which is quantified using the fluorescence of a Ca^2+^ indicator dye (Fig. 1C). To measure LAPD activity, HEK-TM cells were pre-stimulated with NKH477, a water-soluble forskolin analog that activates transmembrane adenylate cyclases (tmACs) and, thereby, increases cAMP levels, leading to a Ca^2+^ influx. NKH477 stimulation was performed under illumination with 850 nm that deactivates LAPD, as previously described [21, 22]. When the Ca^2+^ influx reached a steadystate, LAPD was activated by 690 nm light, decreasing cAMP levels and, thereby, the intracellular Ca^2+^ concentration (Fig. 1C). We measured the mNphp3(201)-bPAC-mCherry or mNphp3(201)-LAPD-mCherry activity and compared it to the non-ciliary tagged bPAC- or LAPD-mCherry proteins. Light stimulation of mNphp3(201)-bPAC-mCherry or bPAC-mCherry expressing HEK-TM cells resulted in a transient Ca^2+^ increase, which was absent in mCherry-expressing control cells (Fig. 1D). Repetitive light-stimulation with different light pulses reliably increased the intracellular Ca^2+^ concentration in mNphp3(201)-bPAC-mCherry or bPAC-mCherry expressing HEK-TM cells (Fig. 1D). Normalized peak amplitudes of the Ca^2+^ signal evoked after the first light pulse were lower in mNphp3(201)-bPAC-mCherry than in bPAC-mCherry expressing HEK-TM cells (Fig. 1E), indicating that the N-terminal fusion to a ciliary targeting sequence interferes with the light-dependent activation of bPAC. Next, responses of HEK-TM cells stably expressing mNphp3(201)-LAPD-mCherry or LAPD-mCherry to NKH477 stimulation were quantified: in both, mNphp3(201)-LAPD-mCherry and LAPD-mCherry expressing HEK-TM cells, NKH477 stimulation induced a Ca^2+^ increase (Fig. 1F). Activating LAPD with 690 nm light significantly decreased the intracellular Ca^2+^ concentration in LAPD-mCherry expressing, but not in mNphp3(201)-LAPD-mCherry expressing HEK-TM cells (Fig.1 G), demonstrating that N-terminal fusion to a ciliary targeting sequence interferes with the light-dependent activation of LAPD. Taken together, our results obtained with two different optogenetic tools revealed that fusion with the minimal ciliary targeting motif mNphp3(201) interfered with their light-dependent activation, thus hampering a direct targeting strategy that does not rely on introducing a functional GPCR to the cilium.

**Figure 1:**
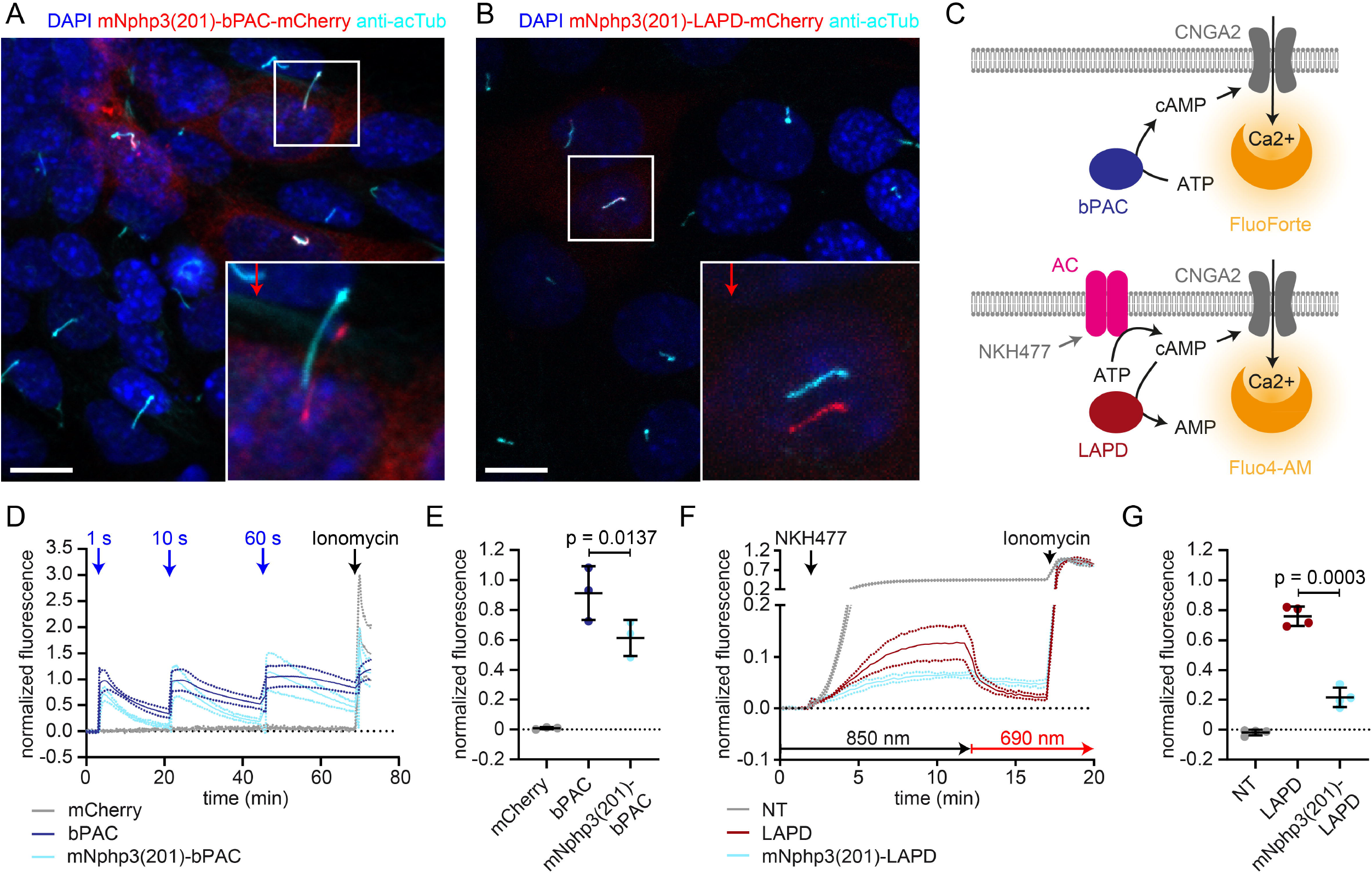
Direct ciliary targeting of optogenetic tools impairs protein function. **A.** Localization of mNphp3(201)-bPAC-mCherry to primary cilia. mIMCD-3 cells expressing mNphp3(201)-bPAC-mCherry were labeled with an anti-acetylated tubulin antibody (cyan, ciliary marker) and with DAPI (blue) to label the DNA. The box indicates the position of the magnified view shown at the bottom right. Red arrow indicates the direction and the length of the shift of the respective fluorescence channel. Scale bar: 10 μm. **B.** Localization of mNphp3(201)-LAPD-mCherry to primary cilia. mIMCD-3 cells expressing mNphp3(201)-LAPD-mCherry were labeled with an anti-acetylated tubulin antibody (cyan, ciliary marker) and DAPI (blue) to label the DNA. The box indicates the position of the magnified view shown at the bottom right. Red arrow indicates the direction and the length of the shift of the respective red channel. Scale bar: 10 μm. **C.** Assays to measure bPAC or LAPD activity using Ca^2+^ imaging. HEK293 cells express the CNGA2-TM ion channel, which opens upon cAMP binding and conducts Ca^2+^ (HEK-TM) [33]. Light-dependent activation of bPAC increases intracellular cAMP levels, leading to a Ca^2+^ influx, which was quantified using a fluorescent Ca^2+^ dye (GFP-certified FluoForte). To measure LAPD activity, HEK-TM cells were pre-stimulated with 100 μM NKH477 to activate transmembrane adenylate cyclases (AC), thus increasing cAMP levels. Ca^2+^ influx was detected by a Ca^2+^ dye (Fluo4-AM). **D.** Quantification of bPAC activity. GFP-certified-FluoForte-loaded HEK-TM cells expressing mCherry only (grey), bPAC-mCherry (blue), or mNphp3(201)-bPAC-mCherry (cyan) were stimulated with 465 nm light pulses (1 mW/cm^2^) of different length and the increase in the intracellular Ca^2+^ concentration was measured. To evoke a maximal Ca^2+^ response, cells were stimulated with 2 μM ionomycin. Data are shown as mean ± SD (dotted lines) for the normalized fluorescence (F-F(baseline))/(F(ionomycin)-F(baseline))/fraction of mCherry-positive cells, n = 3 independent experiments (each data point represents the average of a duplicate or triplicate measurement). **E.** Mean peak amplitudes of the Ca^2+^ signal at 3-6 min after the first light pulse. Data are shown as individual data points and mean ± SD, n = 3. **F.** Quantification of LAPD activity. Fluo4-AM-loaded HEK-TM cells expressing LAPD-mCherry (red) or mNphp3(201)-bPAC-mCherry (cyan) were incubated with 100 μM NKH477 during continuous 850 nm light stimulation (0.5 μW/cm2). At steady-state, light stimulation was switched to 690 nm (0.5 μW/cm2). NT: non-transfected cells (grey). Data are shown as mean ± SD (dotted lines) for the normalized fluorescence (F-F(baseline))/(F(ionomycin)-F(baseline)). **G.** Mean decrease of the Ca^2+^ signal after 690 nm light stimulation (fraction of maximum value after NKH477 increase), determined over 45 s at 3 min after switching to 690 nm. Data are shown as individual data points and mean ± SD, n = 4 independent experiments (each data point represents the average of a duplicate or triplicate measurement); p-values calculated using a paired, two-tailed t-test are indicated. NT: non-transfected cells.

### Targeting optogenetic tools to the primary cilium using nanobodies

We next devised a combinatorial strategy that allows targeting to the primary cilium, while entirely avoiding N-terminal fusion. Rather, we fused our optogenetic tools with a fluorescent reporter (e.g. mCherry) at their C termini, which leaves photoactivation unaffected [19, 22]. To direct these proteins to primary cilia, we co-expressed a nanobody, which is directed against the tag (mCherry) and is fused to the ciliary targeting sequence mNphp3(201) at its N terminus. We hypothesize that the nanobody binds to its target in the cytoplasm, the nanobody-protein-complex is recognized by the ciliary targeting machinery, and is then transported into the primary cilium (Fig. 2A). We first tested the anti-mCherry nanobody VHH_LaM-2_ [34, 35], fused to eGFP at the C terminus and mNphp3(201) at the N terminus, in mIMCD-3 cells. Indeed, the nanobody localized to primary cilia (Fig. 2B). Next, we assessed whether nanobody binding was sufficient to traffic our optogenetic tools to the cilium. Co-expression of the nanobody with LAPD-mCherry resulted in ciliary localization of both the nanobody fusion and LAPD-mCherry (Fig. 2C). In contrast, LAPD-mCherry remained exclusively cytosolic in the absence of the nanobody (Fig. 2D). A second nanobody directed against mCherry, VHH_LaM-4_, [34, 35] also localized to primary cilia and resulted in ciliary localization of LAPD (Sup. Fig. 1A, B), while a nanobody directed against eGFP [36] did not mediate ciliary localization of LAPD-mCherry (Sup. Fig. 1C). The nanobody-based targeting approach also succeeded in localizing bPAC-mCherry to the primary cilium (Fig. 2E). Taken together, nanobody-based targeting of optogenetic toosl was efficient and specific. Hence, we assume that our approach is generally applicable to target proteins of interest to cilia.

**Figure 2:**
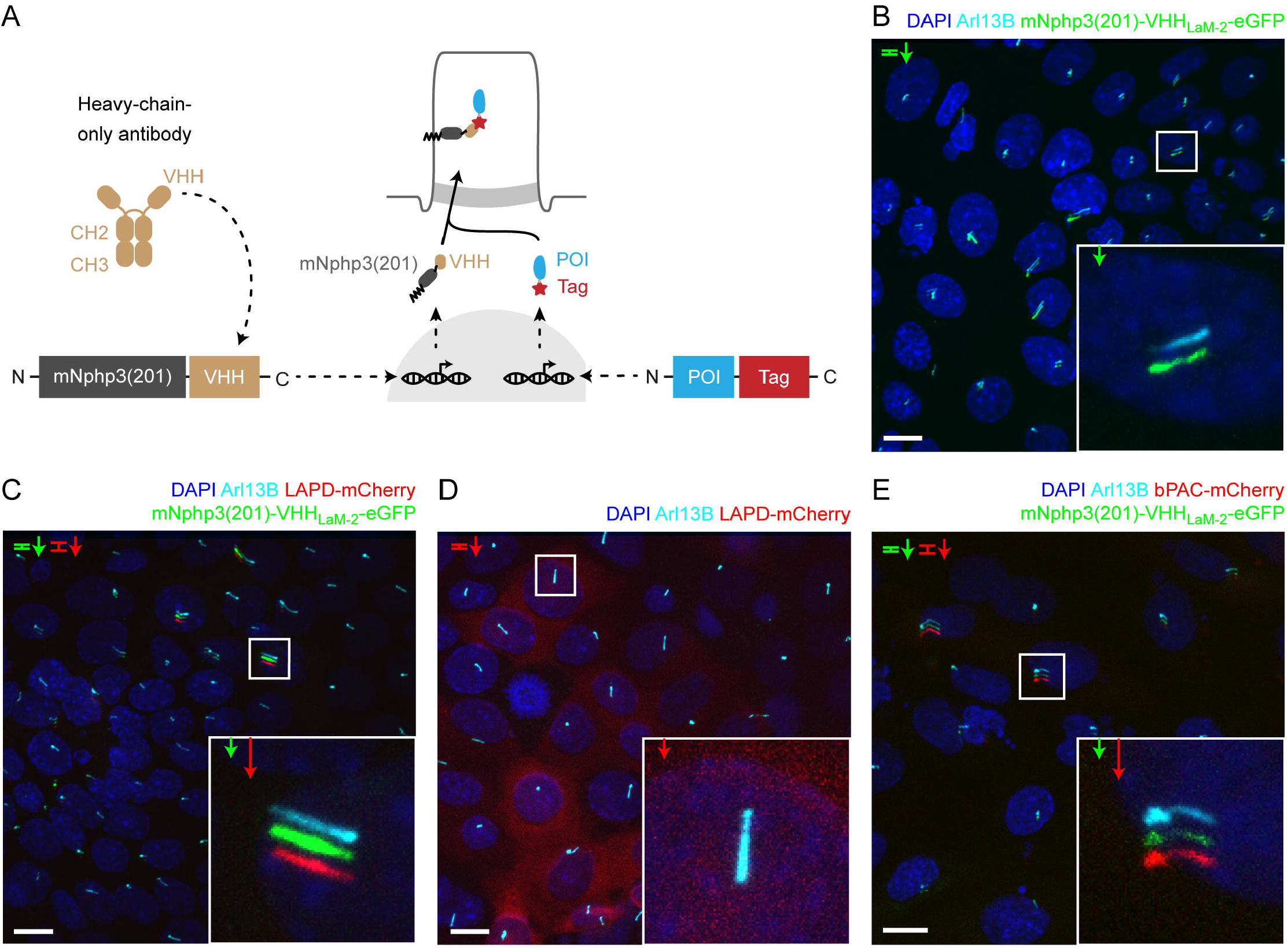
Targeting optogenetic tools to the primary cilium using nanobodies. **A.** Schematic overview targeting approach. Nanobodies were fused to the C terminus of mNphp3(201) for ciliary localization. The protein of interest (POI) is co-expressed with a C-terminal tag or fusion partner that is recognized by the nanobody. Binding of the nanobody to the tag is expected to result in ciliary localization of the POI. **B.** Localization of the anti-mCherry nanobody (VHH_LaM-2_) to primary cilia. mIMCD-3 cells were transfected with mNphp3(201)-VHH_LaM-2_-eGFP (green). **C.** Localization of the anti-mCherry nanobody and LAPD-mCherry to primary cilia. mIMCD-3 cells were co-transfected with mNphp3(201)-VHH_LaM-2_-eGFP (green) and LAPD-mCherry (red). **D.** Cytoplasmic localization of LAPD-mCherry. mIMCD-3 cells were transfected with LAPD-mCherry (red). **E.** Localization of the anti-mCherry nanobody and bPAC-mCherry to primary cilia. mIMCD-3 cells were co-transfected with mNphp3(201)-VHH_LaM-2_-eGFP (green) and bPAC-mCherry (red). All cells shown in B-E were labeled with an Arl13B antibody (cyan, ciliary marker) and DAPI (blue). All scale bars: 10 μm. Boxes indicate the position of the magnified view shown at the bottom right. Arrows in different colors indicate the direction and the length of the shift of the respective fluorescence channel.

### Nanobody binding does not interfere with photoactivation

To test whether nanobody binding interferes with the light-dependent activation of LAPD or bPAC, we first tested their activity in non-ciliated cells to directly compare bound and nonbound optogenetic tools and then verified the optimal experimental condition in ciliated mIMCD-3 cells. To measure the activity in HEK-TM cells, bPAC- or LAPD-Cherry were coexpressed with the cilia-targeted mCherry nanobody mNphp3(201)-VHH_LaM-2_-eGFP. In the absence of the nanobody, bPAC- and LAPD-mCherry displayed a cytosolic distribution (Sup. Fig. 2A, B). In the absence of primary cilia, the mNphp3(201)-VHH_LaM-2_-eGFP nanobody formed clusters in HEK-TM cells (Sup. Fig. 2C), while the VHH_LaM-2_-eGFP nanobody did not (Sup. Fig. 2D). Co-expression of bPAC- or LAPD-mCherry with the mNPHP3(201)-tagged nanobody resulted in cluster localization of either bPAC or LAPD, demonstrating that the nanobody interacts with the mCherry fusion-proteins in the cytoplasm (Sup. Fig. 2E, F).

To test bPAC or LAPD function in the presence of the nanobody, we compared the light-dependent activation of bPAC or LAPD in the presence or absence of the mNPHP3(201)-tagged mCherry nanobodies VHH_LaM-2_ and VHH_LaM-4_ (fused to either eGFP or a hemagglutinin HA-tag, respectively) or in the presence of a ciliary protein that does not interact with either bPAC- or LAPD-mCherry (Sstr3-eGFP). Under each condition, photoactivation of bPAC or LAPD activity was retained, demonstrating that interaction with the nanobody did not interfere with protein function (Sup. Fig. 3A, B).

To scrutinize bPAC and LAPD activity during nanobody-binding in a ciliary context, we aimed to use cAMP biosensors, which can be targeted to the primary cilium. However, all reported biosensors spectrally overlap with the LAPD activation spectrum to some extend [37]. For bPAC, one biosensor is well suited to simultaneously activate bPAC and measure changes in cAMP levels: the red-shifted cAMP biosensor R-FlincA [38]. We co-expressed bPAC-eGFP and R-FlincA and first tested this approach in the cell body (Fig. 3A). Photoactivation of bPAC-eGFP in HEK293 cells transiently increased the R-FlincA fluorescence, whereas a non-binding mutant sensor did not respond to bPAC photoactivation (Fig. 3B, C), demonstrating that a light-stimulated increase in the intracellular cAMP concentration can be concomitantly measured using R-FlincA.

**Figure 3:**
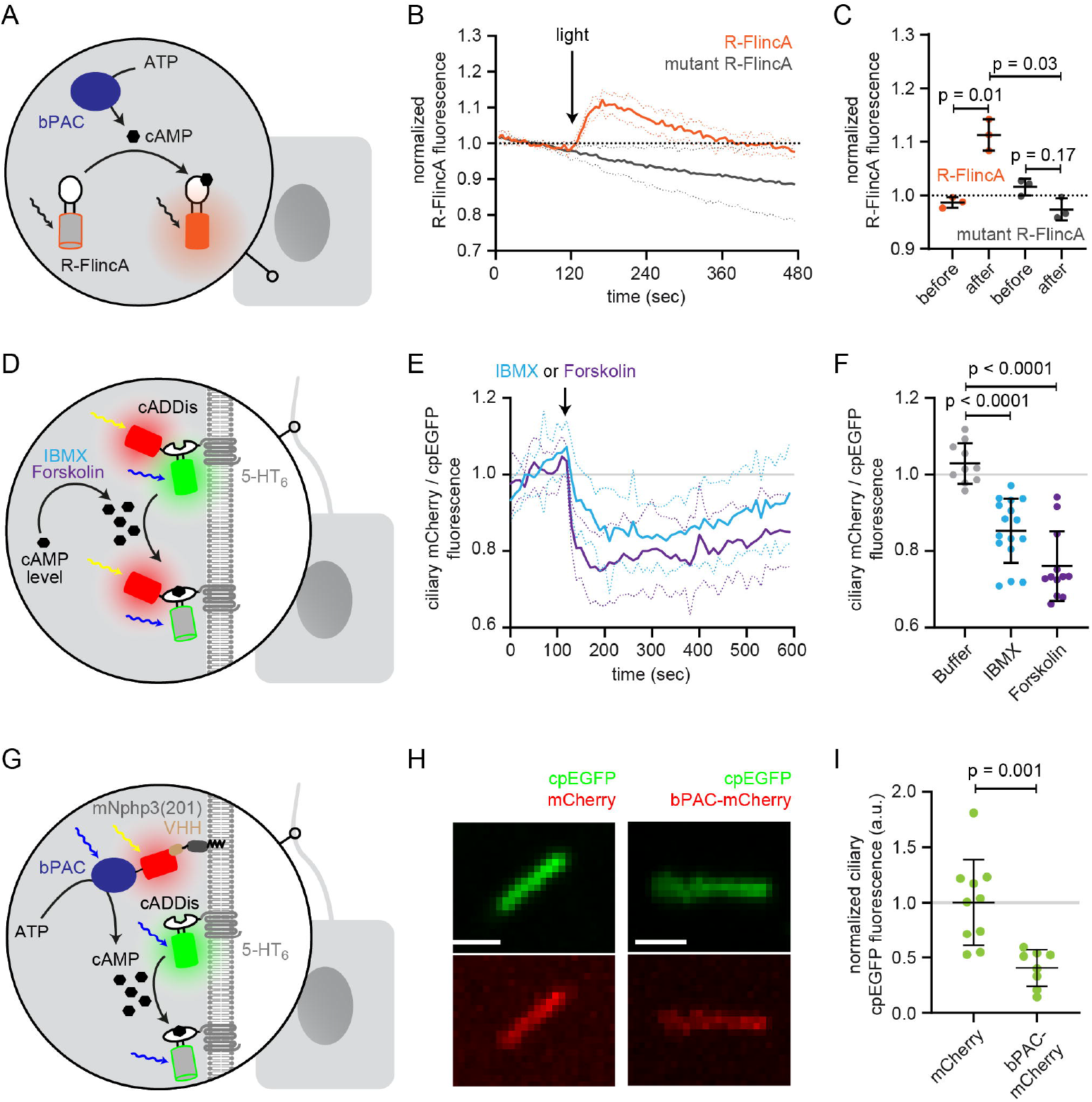
Functional characterization of bPAC in the cell body and cilium. **A.** Schematic overview of the bPAC activity assay in non-ciliated HEK293 cells using R-FlincA (see B-C). **B.** HEK293 cells were transfected with bPAC-eGFP and R-FlincA or the non-binding R-FlincA mutant [38]. The change in R-FlincA fluorescence was measured over time before and after photoactivation of bPAC (5 s, white light, 2.1 mW/cm^2^ at 480 nm). Data are shown as mean (solid lines) ± S.D. (dotted lines), n = 3 with 4 cells per experiment. **C.** Normalized R-FlincA or R-FlincA mutant fluorescence directly before and for the maximal amplitude after photoactivation. Data extracted from C; p-values have been calculated using a paired, twosided Student’s t-test. **D.** Schematic overview of the assay to measure ciliary cAMP dynamics using 5-HT6-mCherry-cADDis after pharmacologically increasing cAMP levels (see E-F). **E.** Ciliary cAMP dynamics measured using 5-HT6-mCherry-cADDis. Cells were stimulated with 250 μM IBMX (light blue) or 40 μM Forskolin (purple). The normalized ratio of ciliary mCherry/cpEGFP fluorescence is shown as mean (solid lines) ± S.D. (dotted lines); p-values have been calculated by paired, two-sided Student’s t-test. **F.** Mean change in the normalized ratio of ciliary mCherry/cpEGFP fluorescence 60-120 s after stimulation with buffer, IBMX, or Forskolin. Data are shown as individual data points, the mean ± S.D. is indicated; p-values have been calculated by a two-sided Mann-Whitney test. **G.** Schematic overview of the assay to measure light-evoked ciliary cAMP dynamics after bPAC stimulation using 5-HT_6_-cADDis (see H-I). **H.** 5-HT6-cADDis fluorescence in cilia with mNphp3(201)-VHH_LaM-2_-HA targeted mCherry or bPAC-mCherry in the first frame of imaging. Scale bar: 2 μm. **I.** Mean normalized ciliary cpEGFP fluorescence in the first frame. All data have been normalized to the mean cpEGFP fluorescence in the mCherry control. Data are shown as individual data points, the mean ± S.D. is indicated; p-values have been calculated by unpaired, two-sided Student’s t-test.

However, we failed to apply this orthogonal system to the primary cilium due to a low signal-to-noise ratio in the cilium. Therefore, we used an alternative approach to measure photoactivation of nanobody-targeted bPAC in the cilium. Mouse IMCD-3 cells were transfected with bPAC-mCherry and mNphp3(201)-VHH_LaM-2_-HA to localize bPAC to the cilium. In addition, cells were transduced with the 5-HT6-cADDis green cAMP biosensor, which is targeted to the primary cilium and reports an increase in cAMP with a decrease in cpEGFP fluorescence [12]. We first verified 5-HT6-cADDis sensor function by pharmacologically increasing cAMP levels (Fig. 3D-F). To this end, we used 40 μM Forskolin, an activator of transmembrane adenylyl cyclases, or 250 μM IBMX, a broad-band phosphodiesterase inhibitor. Of note, pharmacological stimulation increases cAMP levels in the whole cell and not specifically in the primary cilium. The sensor reliably reported an increase in cAMP levels in the primary cilium evoked by either of the two stimuli (Fig. 3D, F). Next, we analyzed cAMP levels in primary cilia in the presence of mNphp3(201)-VHH_LaM-2_-HA and bPAC-mCherry or mCherry as control (Fig. 3G-I). In this approach, however, the tools are not spectrally separated as both bPAC and the cAMP biosensor are activated/excited by blue light (here: 488 nm). Thus, measuring the cpEGP fluorescence of the cADDis sensor concomitantly activates bPAC. Indeed, as soon as the measurement was started, bPAC was activated, cAMP levels increased and, in turn, the cpEGFP fluorescence of the cADDis sensor was significantly lower compared to the cpEGFP fluorescence in cilia that contained mCherry only (Fig. 3H, I), demonstrating that photoactivation of bPAC increased cAMP levels in the cilium. Thus, even though the approach lacks the spectral separation, it confirms light-induced changes of cAMP levels in the cilium and demonstrates that nanobody-based targeting of bPAC-mCherry to the cilium increases ciliary cAMP levels after photoactivation.

In summary, our nanobody-based approach provides a versatile means for ciliary targeting without interfering with protein function.

### Targeting of a genetically-encoded biosensor to the primary cilium

We previously engineered and applied a genetically-encoded biosensor, named mlCNBD-FRET, to measure cAMP dynamics in motile cilia [18]. We already demonstrated targeting of this sensor to primary cilia by fusing it to the C terminus of Sstr3 [18]. However, Sstr3 is a functional GPCR, which may interfere with ciliary signaling, in particular cAMP signaling, upon overexpression. We aimed to optimize the targeting approach by fusing mNphp3(201) to the N terminus of mlCNBD-FRET. While the biosensor localized to the primary cilium (Sup. Fig. 4A), biosensor function was severely impaired as mlCNBD-FRET no longer responded to changes in cAMP levels (Sup. Fig. 4B, C). We thus tested whether the biosensor can be targeted to primary cilia using our nanobody-based approach without interfering with protein function. The mlCNBD-FRET sensor consists of the FRET pair cerulean and citrine [18]. Both fluorescent proteins are recognized by the nanobody VHH_enhancer_ directed against eGFP [36, 39]. Fusion of mNphp3(201) to the N terminus of the anti-eGFP nanobody also resulted in ciliary localization (Fig. 4A). In the absence of the nanobody, mlCNBD-FRET was uniformly distributed throughout the cytosol, whereas co-expression with the mNphp3(201)-tagged nanobody resulted in ciliary localization of mlCNBD-FRET (Fig. 4B, C). To test whether nanobody interaction impaired mlCNBD-FRET function, we performed FRET imaging in HEK293 cells expressing mlCNBD-FRET in the presence or absence of the nanobody. Similar to the anti-mCherry nanobody, the cilia-targeted eGFP mNphp3(201)-VHH_enhancer_-mCherry nanobody showed a more clustered subcellular localization in HEK293 cells in the absence of primary cilia formation (Sup. Fig. 4D). Consistently, when binding to the nanobody, mlCNBD-FRET also formed clusters within the cytosol (Sup. Fig. 4E), which did not occur in the presence of mCherry only (Sup. Fig. 4F). To functionally test the FRET sensor in the presence of the nanobody, we first assessed the impact of the nanobody on the fluorescence intensity of the two fluorophores, cerulean and citrine. HEK293 cells were transfected with cerulean or citrine and the eGFP VHH_enhancer_-mCherry nanobody or mCherry only. The fluorescence intensity of cerulean or citrine was normalized to the mCherry fluorescence in the same cell. Both cerulean and citrine showed an increase in fluorescence in the presence of the nanobody compared to the mCherry control as previously described [36], but the relative change for each of the fluorophores was not substantially different (Sup. Fig. 4G). To test whether mlCNBD-FRET:nanobody complexes still respond to changes in cAMP levels, we first measured cAMP-induced FRET changes in non-ciliated HEK293 cells and then in ciliated mIMCD-3 cells. To increase the intracellular cAMP concentration, cells were stimulated with 20 μM isoproterenol, which stimulates AC activity through signaling via GPCRs (G protein-coupled receptors). We analyzed FRET changes in HEK293 mlCNBD-FRET cells co-expressing mNphp3(201)-VHH_enhancer_-mCherry or the non-targeted VHH_enhancer_-mCherry nanobody (Fig. 4D). In the presence of the VHH_enhancer_-mCherry nanobody, the FRET response to stimulation with isoproterenol remained unchanged (Fig. 4E, F) and also interaction with the mNphp3(201)-tagged nanobody only marginally reduced the FRET response and generally left the reporter functional (Fig. 4E, F).

**Figure 4:**
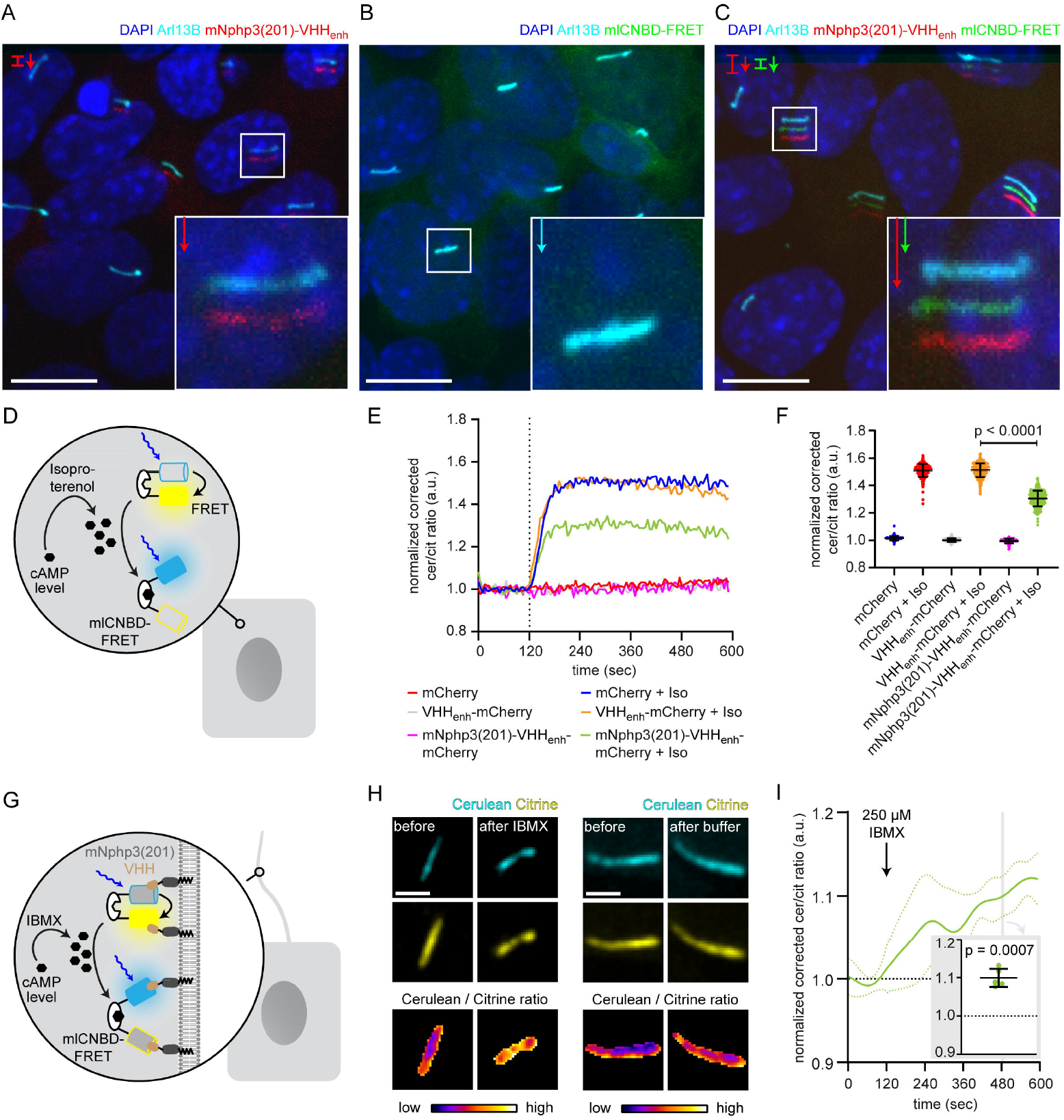
Functional characterization of nanobody-targeted cAMP biosensor. **A**. Localization of the mNphp3(201)-VHH_enhancer_-mcherry anti-eGFP nanobody to primary cilia. **B./C.** Localization of mlCNBD-FRET in mIMCD-3 cells in the **B.** absence or **C.** presence of mNPHP3(201)-VHH_enhancer_-mCherry. **D.** Schematic overview of mlCNBD-FRET imaging in non-ciliated HEK293 cells (see E-F). **E.** FRET imaging in HEK293 mlCNBD-FRET cells transiently co-expressing mCherry, VHH_enhancer_-mCherry, or mNphp3(201)-VHH_enhancer_-mCherry under control conditions or after stimulation with 20 μM isoproterenol (Iso, addition depicted with dotted line). Data are shown as mean (n = 3 independent experiments, 30-90 cells per experiment). **F.** Comparison of maximal change for data shown in E. Data are presented as individual data points and mean ± S.D.; p-value calculated using an unpaired, two-tailed Mann-Whitney test is indicated. **G.** Schematic overview of mlCNBD-FRET imaging in the primary cilium of mIMCD-3 cells (see H-I). **H.** FRET imaging in primary cilia of mIMCD-3 cells expressing mlCNBD-FRET and mNphp3(201)-VHH_enhancer_-mCherry. Cells have been stimulated with 250 μm IBMX (left) or buffer only (right). Cerulean and citrine are shown before and after stimulation with IMBX. The change in cerulean/citrine ratio is shown below (colorscheme indicated at the bottom). Scale bar: 2 μm. **I.** Time course of mean change in FRET (dark green line) ± S.D. (dotted green line) for data set, exemplary shown in H; n = 5. Inset: each data point shows the time-average per cilium at the position indicated by grey box; one-sample Student’s t-test compared to 1.0 indicated.

After having verified biosensor function in the presence of the nanobody in non-ciliated cells, we performed FRET imaging in cilia of mIMCD-3 cells co-expressing mlCNBD-FRET and mNphp3(201)-VHH_enhancer_-mCherry (Fig. 4G). In response to stimulation with 250 μM IBMX to increase cAMP levels, the ciliary-localized mlCNBD-FRET responded with a change in FRET, whereas buffer addition did not change FRET (Fig. 4H, I), demonstrating that the nanobody-targeted mlCNBD-FRET sensor can be used to study cAMP dynamics in the primary cilium. In conclusion, the nanobody-based approach applies not only for targeting optogenetic tools, but also genetically-encoded biosensors to the primary cilium.

### Applying the nanobody-based ciliary targeting approach *in vivo*

Having shown that our bipartite strategy for localization to primary cilia works *in vitro*, we wondered whether we could also target proteins of interest to primary cilia *in vivo*. We first confirmed the ciliary localization of the nanobody *in vivo* by injecting mRNA of the anti-mCherry mNphp3(201)-VHH_LaM-2_-eGFP nanobody into nacre (*mitfa*^-/-^) zebrafish embryos, which are transparent and, therefore, widely used for fluorescence imaging [40]. The cilia-targeted nanobody was expressed and localized to cilia in all tissues, including the developing neural tube, the primary and motile cilia of the spinal cord [41], and the eye (Fig. 5A, B, C) and allowed to mark cilia in an *in vivo* imaging approach (Supp. Fig. 5A). Localization of the nanobody to cilia is similar to the previously described *bactin:arl13b-gfp* transgenic line, where GFP is fused to the ciliary Arl13B protein (Supp. Fig. 5B) [42, 43]. To test whether the cilia-targeted nanobody can also direct proteins to primary cilia *in vivo*, we injected mRNA of the anti-mCherry nanobody fusion mNphp3(201)-VHH_LaM-2_-eGFP into transgenic zebrafish embryos ubiquitously expressing RFP (*Ubi:zebrabow*) [44], which is also bound by the anti-mCherry nanobody. In the absence of the nanobody, RFP was distributed in the cytosol (Fig. 5D, Supp. Fig. 5C). In the presence of the nanobody, RFP was highly enriched in primary cilia (Fig. 5E, Supp. Fig. 5D), demonstrating that our nanobody-based approach efficiently targets ectopically expressed proteins to primary cilia and to motile cilia *in vitro* and *in vivo*.

**Figure 5:**
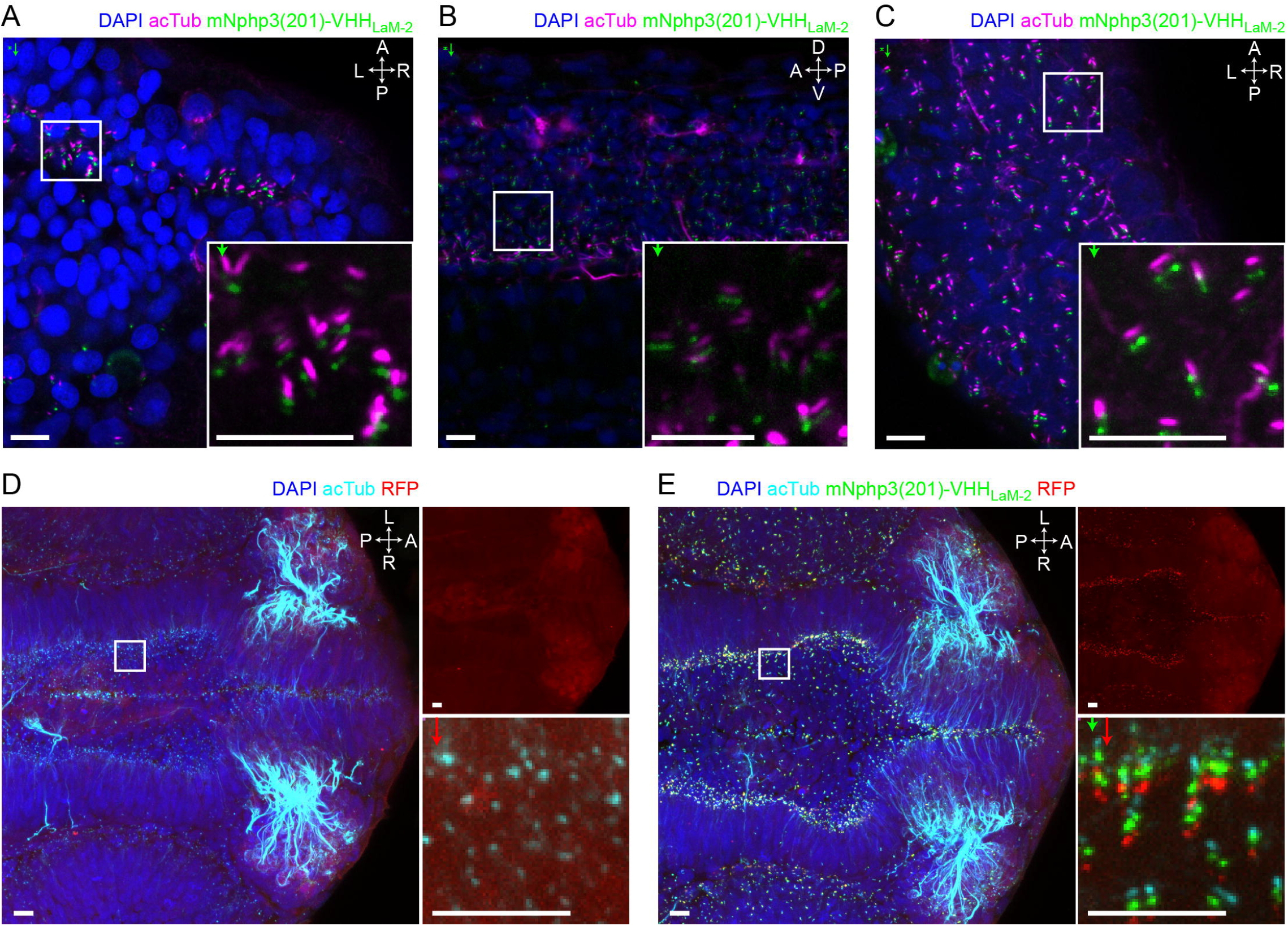
Nanobody-based ciliary protein targeting *in vivo*. **A.** Nanobody localization in the neural tube of a zebrafish embryo. The mRNA of the anti-mCherry mNphp3(201)-VHH_Lam-2_-eGFP nanobody was injected into nacre zebrafish embryos. Embryos were stained with an anti-acetylated tubulin antibody (magenta, ciliary marker), an anti-GFP antibody (green), and DAPI (blue). **B.** see A. for spinal cord. **C.** see A. for eye. **D.** RFP (red) expression in the neural tube of Ubi:zebrabow (Pan et al., 2013) transgenic embryos. **E.** RFP (red) expression in the neural tube of Ubi:zebrabow (Pan et al., 2013) transgenic embryos, injected with mRNA of the anti-mCherry mNphp3(201)-VHH_LaM-2_-eGFP nanobody. Scale bars: 20 μm, magnified view: 10 μm. Boxes indicate the position of the magnified views shown at the bottom right as inset (A-C) or as a separate panel next to the overview image (D, E). Arrows in different colors indicate the direction and the length of the shift of the respective fluorescence channel. The upper right panel in D and E shows the RFP channel only, the bottom right panel shows the magnified view. A: anterior, P: posterior, L: left, R: right, D: dorsal, V: ventral. All images were taken from fixed samples.

### Investigating the spatial contribution of cAMP signaling to cilia length control

The primary cilium is a dynamic cellular structure that assembles and dissembles in accordance with the cell cycle [45–47]. The interplay between assembly and disassembly determines the length of the primary cilium. cAMP-dependent signaling pathways have been shown to regulate cilia length [14–16, 48, 49]. Changes in cAMP signaling to study cilia length control have only been evoked using pharmacology, lacking spatial resolution and targeting both, the cilium and cell body. However, it is generally accepted that cAMP signaling occurs within defined subcellular compartments to evoke a specific cellular response [50]. Whether an increase in cAMP levels in either the cilium or the cell body is sufficient to evoke a change in ciliary length, is not known. Thus, it is not surprising that it has been controversially discussed whether an increase in the intracellular cAMP concentration, evoked by pharmacological stimulation, results in an increase or decrease in cilia length, [14, 15], as spatial cAMP signaling might evoke a differential response, which is impossible to reveal using pharmacology. We also performed pharmacological stimulation of cAMP synthesis in mIMCD-3 cells using Forskolin and analyzed the change in cilia length. To analyze cilia length in an automated and unbiased fashion in 3D, we developed an ImageJ plug-in called CiliaQ. In fact, the response was quite variable and we did not observe a significant change in cilia length (Fig. 6A). Thus, we set out to investigate the spatial contribution of an increase in cAMP levels in the cilium or cell body in regulating cilia length.

**Figure 6:**
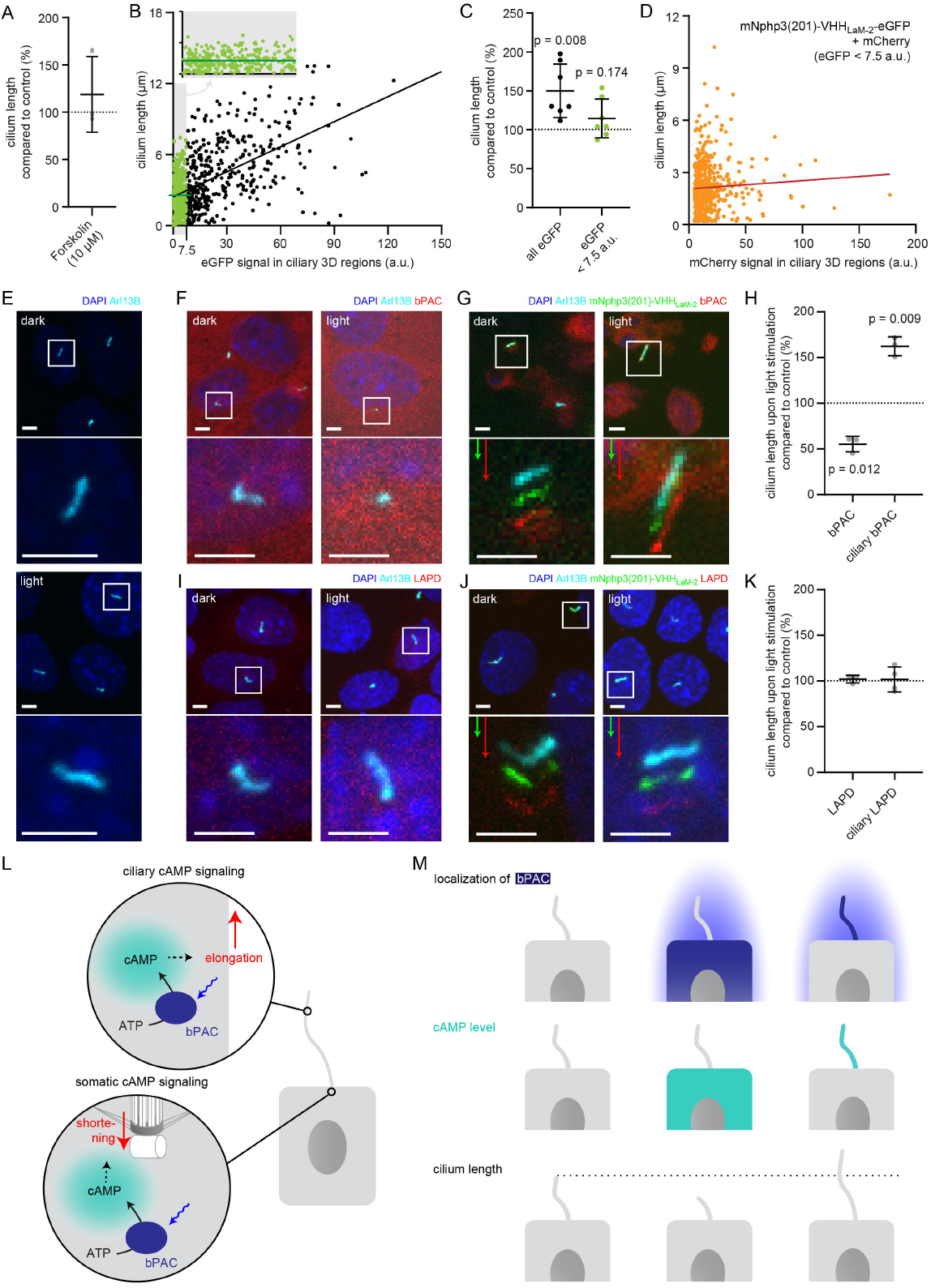
Controlling cilia length using optogenetics. **A.** Cilia length of mIMCD-3 cells stimulated for 1 h with 10 μM Forskolin (solvent: DMSO), normalized to the DMSO control. Data are shown as mean ± S.D., n = 3 with at least 40 cells per experiment. **B.** Correlation of cilia length and eGFP fluorescence (a.u., average ciliary fluorescence of non-transfected control cells was subtracted) in the cilium in mIMCD-3 cells transiently expressing mNphp3(201)-VHH_Lam-2_-eGFP. Below 7.5 a.u., the cilia length is independent of the eGFP fluorescence (see inset, values are highlighted in green, slope not different from zero, correlation: p = 0.07), whereas including values > 7.5 a.u., there is a linear correlation between the cilia length and the eGFP fluorescence in the cilium (slope different from zero, correlation: p < 0.0001). **C.** Length of cilia that show mNphp3(201)-VHH_LaM-2_-eGFP localization and an eGFP fluorescence < 7.5 a.u., normalized to equally treated, non-transfected (NT) control cells. Data are shown as mean ± S.D., n = 7 with at least 18 cilia per experiment; p-values determined using unpaired, two-tailed Student’s t-test are indicated. **D.** Correlation of cilia length and mCherry fluorescence in the cilium in mIMCD-3 cells transiently expressing mNphp3(201)-VHHLaM-2-eGFP and mCherry. Only cilia with an eGFP fluorescence below 7.5 a.u. were taken into account. There is no linear correlation between the mCherry fluorescence and cilia length (slope not different from zero, correlation: p = 0.2). **E.** mIMCD-3 cells (non-transfected, NT) kept in the dark (top) or stimulated with light (bottom, 1 h, 465 nm, 38.8 μW/cm2) **F.** mIMCD-3 bPAC-mCherry cells kept in the dark (left) or stimulated with light (right, 16 h, 465 nm, 38.8 μW/cm2). **G.** mIMCD-3 bPAC-mCherry transiently transfected with mNphp3(201)-VHH_LaM-2_-eGFP kept in the dark (left) or stimulated with light (right, 1 h, 465 nm, 38.8 μW/cm2). **H.** Normalized cilia length after light stimulation (left 1 h, right 16 h; 465 nm, 38.8 μW/cm2) for mIMCD-3 bPAC-mCherry cells with or without transiently expressing mNphp3(201)-VHH_LaM-2_-eGFP. Only cilia with an eGFP fluorescence < 7.5 a.u. were included and each data point was normalized to control cells. Data are shown as mean ± S.D., n = 3 with at least 25 cells per experiment; p-values determined using one-sample Student’s t-test compared to 100% are indicated. I. mIMCD-3 LAPD-mCherry cells kept in the dark (left) or stimulated with light (right, 16 h, 630 nm, 42.3 μW/cm2). J. mIMCD-3 LAPD-mCherry transiently transfected with mNphp3(201)-VHH_Lam-2_-eGFP kept in the dark (left) or stimulated with light (right, 16 h, 630 nm, 42.3 μW/cm2). K. Normalized cilia length after light stimulation (16 h, 630 nm, 42.3 μW/cm2) for mIMCD-3 LAPD-mCherry with or without transiently expressing mNphp3(201)-VHH_Lam-2_-eGFP. Only cilia with an eGFP fluorescence < 7.5 a.u. were included and each data point was normalized to control cells. Data are shown as mean ± S.D., n = 3-4 with at least 18 cells per experiment; p-values determined using one-sample Student’s t-test compared to 100% are indicated. Cells in E-G and I-J were stained with an Arl13B antibody (cyan) and DAPI (blue). All boxes indicate the magnified view below. Arrows indicate the direction and the length of the shift of the respective same-colored fluorescence channel. Scale bar for all images: 3 μm. L. Spatial cAMP signaling controlling cilia length. Our data suggest a model, in which cAMP signaling in the cell body, stimulated by photoactivation of bPAC and an increase in cAMP levels, causes primary cilia shortening, whereas an increase of cAMP levels in the cilium results in primary cilia elongation. M. Summary of the correlation between bPAC localization and photoactivation, cAMP levels, and cilia length.

To this end, we used a monoclonal IMCD-3 cell line stably expressing bPAC-mCherry in combination with the mNphp3(201)-VHH_Lam-2_-eGFP nanobody. Since ectopic expression of a ciliary protein may result in an increase of the cilia length [51, 52], we first tested whether expression of the mNphp3(201)-tagged nanobody in the cilium had an impact on the length of the cilium. Indeed, ectopic expression of the mNphp3(201)-tagged nanobody resulted in longer cilia compared to non-transfected control cells. There was a linear correlation between the expression level of the mNphp3(201)-tagged nanobody in the cilium and cilia length: the higher the expression (determined by eGFP fluorescence), the longer the cilia (Fig. 6B), as has been reported previously for ectopic expression of membrane proteins in the cilium [51]. However, in the low expression regime, i.e. < 7.5 a.u. eGFP fluorescence, there was no linear correlation between the expression level of the mNphp3(201)-tagged nanobody in the cilium and cilia length (Fig. 6B inset, Fig. 6C), demonstrating that ectopic expression of the mNphp3(201)-tagged nanobody in the cilium at a low level does not change ciliary length. To verify whether the mNphp3(201)-tagged nanobody also does not alter cilia length in the low mNphp3(201)-VHH_Lam-2_-eGFP expression regimes (< 7.5 a.u.) while in complex with its target, we analyzed mIMCD-3 cells co-expressing mNphp3(201)-VHH_Lam-2_-eGFP and mCherry. In the low mNphp3(201)-VHH_Lam-2_-eGFP expression regimes (< 7.5 a.u.), there was no linear correlation between the mCherry fluorescence and cilia length (Fig. 6D), demonstrating that targeting the mNphp3(201)-tagged nanobody in complex with another protein to the cilium does not alter cilia length. Our results underline that the amount of protein in the cilium has to be carefully titrated and a thorough analysis is needed to rule out any unspecific effects caused by ectopic expression of ciliary proteins. In the following, we hence only analyzed cilia with a mNphp3(201)-VHH_Lam-2_-eGFP expression level of < 7.5 a.u.. We compared the change in cilia length upon photoactivation of bPAC-mCherry either in the cell body or in the presence of mNphp3(201)-VHH_LaM-2_-eGFP in the cilium (Fig. 6E-H). No light-dependent increase in ciliary length was observed in non-transfected cells (Fig. 6E). Stimulating cAMP synthesis by light in the cell body significantly reduced cilia length (Fig. 6F, H), whereas stimulating cAMP synthesis in the cilium significantly increased cilia length (Fig. 6G, H).

Next, we complemented our analysis by investigating whether reducing cAMP levels in either the cell body or the cilium using photoactivation of LAPD also changed cilia length. We compared the change in cilia length upon photoactivation of LAPD-mCherry either in the cell body or, in the presence of mNphp3(201)-VHH_Lam-2_-eGFP, in the cilium. However, neither stimulation of LAPD activity in the cell body nor in the cilium had an effect on cilia length (Fig. 6I-K), although photoactivation of LAPD reduced basal cAMP levels (Sup. Fig. 6A). Thus, the cAMP-dependent signaling pathways that control the length of the cilium seem to be sensitive to an increase, but not to a decrease of the basal cAMP concentration. In summary, cAMP-dependent signaling pathways in the cell body or the cilium evoke opposing effects on cilia length control, demonstrating how compartmentalized cAMP signaling determines specific ciliary and, thereby, cellular functions (Fig. 6L, M).

## Discussion

The primary cilium constitutes a unique subcellular compartment. However, our understanding of how specific ciliary signaling pathways control cellular functions is still limited. Optogenetics is an apt method to measure and manipulate ciliary signaling independent of the rest of the cell. We have developed a nanobody-based approach to target optogenetic tools or genetically-encoded biosensors to primary cilia *in vitro* and *in vivo*, while preserving protein function. This novel strategy principally extends to other proteins of interest and subcellular domains. The only requirement is fusion of the protein of interest to a tag or partner that is recognized by the nanobody, without impairing protein function. This is combined with fusion of the nanobody to a targeting sequence for the specific subcellular compartment. This approach is particularly useful for proteins, whose function is impaired by fusion to a targeting sequence, as we have shown here for the light-activated phosphodiesterase LAPD and the mlCNBD-FRET sensor.

Optogenetic tools have been used in motile cilia, i.e. sperm flagella [17, 19], where no specific targeting sequence is needed. When expressed under the control of the sperm-specific protamine-1 promoter, bPAC exclusively localized to sperm flagella and allowed to control sperm signaling and function by light [19]. However, this approach does not work for primary cilia. Genetically-encoded biosensors for cAMP and Ca^2+^ have already been targeted to primary cilia [12, 18, 53–55]. These sensors have been fused to the C terminus of full-length GPCRs, e.g. 5-HT6 (5-hydroxytryptamine receptor 6) [12]. Yet, overexpression of ciliary proteins like 5-HT6 or Arl13b caused abnormal cilia growth [51, 55], limiting this targeting approach. Our results demonstrate that also high expression levels of the nanobody in the cilium increased cilia length (Fig. 6B). Thus, the amount of protein expressed in the cilium has to be carefully titrated and a thorough analysis is needed to rule out any unspecific effects caused by ectopic expression of the protein and reveal the specific contribution of a protein of interest to ciliary signaling and function.

Targeting proteins to a specific location using nanobodies has been applied to study proteinprotein interaction in cells [56] and *Drosophila* [57]. Herce et al. fused an anti-GFP nanobody to the lac repressor to localize the nanobody and its GFP-fused interaction partner to the nucleus. Using this approach, the authors have studied binding and disruption of p53 and HDM2 (human double minute 2), one of the most important protein interactions in cancer research [56]. Harmansa et al. have mislocalized transmembrane proteins, cytosolic proteins, and morphogens in *Drosophila* to study the role of correct protein localization for development *in vivo* [57]. Targeting to organelles using nanobodies has also been achieved for two specific proteins, p53 and survivin [58, 59]. For primary cilia, recombinantly expressed fusion proteins fused to nanobodies have been used to investigate the ciliary diffusion barrier by diffusion-to-capture assays [60], and determined a free diffusion across the ciliary barrier only for proteins < 100 kDa [61]. Although larger proteins can enter cilia by active transport processes, this size cutoff may limit the size of proteins that can be targeted to cilia by a piggyback mechanism using a cilia-localized nanobody. Importantly, a recently developed nanobody, targeting a small alpha-helical epitope tag of 13 amino acids, NbALFA, also works in the cytosol and may reduce the overall size of the protein complexes targeted to primary cilia [62].

Since several proteins proposed to function in cilia also have well-known functions outside the cilium (such as PKA or AMPK), it is a major challenge in cilia biology to specifically interfere with their cilium-specific functions. One approach is to target proteins or peptides with inhibitory functions specifically to the cilium, e.g. to interfere with PKA activity inside cilia [30]. Cilia length is an important parameter that determines cilia function [63]. However, the spatial contribution of signaling pathways in the cilium or cell body that regulate cilia length is not well understood. So far, only pharmacology that, however, lacks spatial resolution has been used to investigate the role of cAMP in the regulation of cilia length. Using our approach to optogenetically manipulate cAMP levels in the cilium or cell body allows to address the spatial contribution of cAMP signaling in cilia length control. Our results indicate that an increase in cAMP levels and, thereby, cAMP signaling in the cilium or the cell body exerts opposing effects by either increasing or decreasing cilia length, respectively (Fig. 6H). Indeed, it has been demonstrated in HEK293 cells that an increase in cellular cAMP levels and concomitant PKA-dependent protein phosphorylation at centriolar satellites induced ubiquitination and proteolysis of NEK10 by the co-assembled E3 ligase CHIP and promoted cilia resorption [15]. This is supported by elegant studies of cilia length control in *Chlamydomonas* [47, 64–72]. Here, the IFT machinery and its regulation by phosphorylation plays an important role. There is a negative correlation between IFT particles entering flagella and flagella length, suggesting a length-dependent feedback control of IFT entry and, thereby, flagella length. At least in *Chlamydomonas*, phosphorylation of the kinesin-II subunit Fla8 (pFla8) by the calcium-dependent kinase 1 (CDPK1), a homolog of CAMKII, changes the rate of IFT entry and, thereby, flagella length [66]. Thus, a cellular sensing system controls pFla8 levels, reduces the rate of IFT entry and controls flagella length. In contrast, stimulation of cAMP signaling in the cilium results in an increase in cilia length, probably through cAMP/PKA-dependent signaling pathways, as suggested by experiments in primary cilia of epithelia and mesenchymal cells, where cilia elongation is induced by a cAMP/PKA-dependent mechanism [14]. In line with this finding, an increase in intracellular cAMP levels and downstream activation of PKA has been shown to contribute to an increase in cilia length by regulating anterograde IFT transport velocity [14].

Whereas a light-stimulated increase in ciliary cAMP levels by bPAC resulted in an increase in cilia length, a decrease in ciliary cAMP levels evoked by photoactivation of LAPD did not change cilia length. Only a cAMP increase in the cell body resulted in a reduction of the cilia length. Thus, cilia length control only seems to be sensitive to an increase, but not to a decrease in cAMP levels, and the direction of the regulation (lengthening or shortening of cilia) appears to be dependent on the origin of the cAMP increase. A possible explanation for this surprising finding is that PKA is the major target for cAMP and has also been implicated in the cAMP-dependent regulation of cilia length control. *In vitro*, PKA is half-maximally activated at 100-300 nM. However, a recent report demonstrated that in fact, the sensitivity of PKA for cAMP is almost twenty times lower in cells compared to *in vitro* [52]. In many different cell types, resting cAMP levels have been shown to be around 1 μM [52], and primary cilia in mIMCD-3 cells contain a similar cAMP concentration as the cell body [55]. Based on these numbers, we hypothesize that cilia length is mainly regulated by PKA-dependent mechanisms, only responding to an increase of the local cAMP concentration. As described above, this increase has to be well above > 1 μM to activate PKA in cells. In turn, reducing basal cAMP levels below 1 μM by photoactivation of LAPD would not result in a change in ciliary length since the activity of PKA, compared to basal levels, would not be changed.

Our approach and complementary developments in other labs using nanobodies for subcellular targeting [56–59] open up new avenues for analyzing signaling pathways in primary and motile cilia, as demonstrated by the application of the cilia-targeted nanobody in zebrafish, and beyond that in many other subcellular domains *in vitro* and *in vivo*. The nanobody-based targeting approach conveys high specificity through a strong interaction with its binding partner (Kd ~ 1 nM), which is a prerequisite for subcellular targeting and binding of endogenously expressed proteins *in vitro* and *in vivo* [73, 74]. In addition, nanobodies are small (~ 15 kDa) and genetically-encoded on one single open reading frame [73]. Thus, the generation of transgenic animals, e.g. mice or zebrafish, expressing the nanobody targeted to a subcellular compartment is straightforward. Analogous to the Cre/loxP recombinase system, the nanobody-based targeting approach offers endless combinations of targeting any protein of interest to a desired subcellular compartment by simply crossing different transgenic lines. This will greatly facilitate the analysis of cellular signaling in the whole cell and in a specific compartment *in vivo*. Conditional expression patterns will further allow temporally controlled recruitment or cell-type specific localization. Many transgenic animals have already been generated, expressing optogenetic tools that are either fused to a fluorescent protein or even contain fluorescent proteins as functional read-out, e.g. biosensors for second messengers. Combining these transgenic lines with transgenic animals that allow nanobody-based subcellular targeting will pave the way for a more systematic and nuanced application of optogenetics to study cellular signaling. Such studies will unravel how subcellular compartments control cellular functions under physiological and pathological conditions.

## Acknowledgement

The project was supported by grants from the Deutsche Forschungsgemeinschaft (DFG): SPP1926: grant MO2192/4-1 (to AM) and grant WA3382/2-1 (to DW), SPP1726: grant WA3382/3-1 (to DW), TRR83/SFB (to DW), FOR2743 (to DW), and under Germany’s Excellence Strategy – EXC2151 – 390873048 (to DW and FIS), Emmy Noether (to FIS), the Boehringer Ingelheim Fonds (to JNH), and a Samarbeidsorganet Helse Midt-Norge grant (to NJY). We thank Jens-Henning Krause for technical support, the Core Facility Nanobodies of the University of Bonn, the Microscopy Core Research Facility of the Bonn Technology Campus, the Core Research Facility for Light Microscopy (CRFS) of the DZNE (German Center for Degenerative Diseases), and the fish facility support team at the Kavli Institute for Systems Neuroscience.

## Author contribution

DW, JNH, AM, NJY, DUM, and FIS designed the experiments and wrote the manuscript. WB performed the cloning, JNH, RC, FK, NJY, BS, and CV performed the experiments. DW, AM, and FIS acquired the funding to conduct the experiments.

## Materials and methods

### Plasmids

Coding sequences for codon-optimized anti-mCherry nanobodies VHH_LaM-2_ and VHH_LaM-4_ were synthesized as GeneBlocks by IDT, based on the amino acid sequences provided by Fridy et al., 2018. A vector encoding the anti-eGFP nanobody GBP-1 (VHH_Enhancer_) [36] was kindly provided by the lab of Hidde Ploegh (Boston Children’s Hospital, Boston, MA, USA). Information regarding the cloning can be found in the Supplementary Information and in Supplementary Table 1.

### Cell lines and tissue culture

HEK293 (CRL-1573) and mIMCD-3 (CRL-2123) cells were obtained from American Type Culture Collection (ATCC). HEK293 TM (HEK-TM) cells were generated as described previously [33]. HEK-TM cells were transfected with pc3.1-bPAC-mCherry or pcDNA6-LAPD-mCherry and selected for stable expression. HEK-mlCNBD-FRET were generated as described previously [18]. Information regarding the cultivation of the cell lines can be found in the Supplementary Information.

### Transfection

mIMCD-3 cellls were transfected with Lipofectamine 2000 (Thermo Fisher Scientific) and HEK293 cells with polyethylenimine (PEI, Sigma Aldrich). Details are described in the SI Appendix. All cells referred to as non-transfected (NT) were subjected to the same transfection protocol as transfected cells, but without adding DNA, Lipofectamine and PEI.

### Immunocytochemistry

Immunocytochemistry was performed according to standard protocols. Detailed information can be found in the Supplementary Information.

### Optogenetic stimulation for cilia length measurements

mIMCD-3 cells and mIMCD-3 bPAC cells were seeded, transfected with pcDNA3.1-mNphp3(201)-VHH_LaM-2_-eGFP, and induced to form cilia as described above. Cells were kept in the dark during the entire experiment and handled only under dim red (bPAC) or green (LAPD) light, preventing bPAC- or LAPD-activation, respectively. For “light” stimulation, cells were placed on a LED plate (bPAC: 465 nm, 38.8 μW/cm^2^; LAPD: 630 nm, 42.3 μW/cm^2^) for the last 16 h before harvesting or for 1 h at 24 h prior to harvesting (as indicated in the figure legends). Cells were fixed and further analyzed by immunocytochemistry and confocal microscopy.

### Confocal microscopy and image analysis

Confocal z-stacks (step size 0.4-0.5 μm, 60x objective) were recorded with a confocal microscope (Eclipse Ti, Nikon or Olympus FV100). All depicted images show a maximum projection of a z-stack unless differently stated in the figure legend. Information regarding image analysis can be found in the Supplementary Information.

### Ca^2+^ imaging

Ca^2+^ imaging in 96-well plates in a fluorescence plate reader was performed as previously described [19, 22]. A detailed protocol and all changes are described in the Supplementary Information.

### FRET imaging

FRET imaging was performed as previously described [18]. A detailed protocol and all changes are described in the Supplementary Information.

### R-FlincA imaging

HEK293 cells were seeded and transfected with pcDNA4HMB_R-FlincA or pcDNA4HMB_R-FlincAmut (R221E, R335E) [38] (generously provided by Kazuki Horikawa, Tokushima University, Japan) and pEGFP-N1-bPAC (see Supp. Table 1) as described for LAPD activity measurements. Imaging was performed using the CellR Imaging System (Olympus). The experimental recordings were as follows: R-FlincA signal (57% light intensity, 200 ms exposure time, 572/25 excitation filter, mCherry-B-0MF Semrock dichroic mirror, 630/20 emission filter) was measured every 5 s. At 120 s, cells were illuminated for 5 s with 2.1 mW/cm^2^ white light, followed by further recording of R-FlincA signal every 5 s for 480 s. Subsequently, bPAC-GFP fluorescence (12 % light intensity, 100 ms exposure time, 500/20 excitation filter, M2CFPYFP dichroic mirror, 535-30 emission filter) was measured. Data were analyzed using Fiji/ImageJ (ImageJ Version 1.52i) by selecting low-expressing bPAC-GFP cells with freehand ROIs and determining the mean fluorescence intensity for each ROI in the average signal recorded during the 120 s before white light exposure. Data were plotted as a change of fluorescence over time. Data were acquired from n = 3 measurements.

### Imaging of primary cilia

mIMCD-3 cells were seeded on PLL (0.1 mg/ml, Sigma Aldrich)-coated chambers (μ-Slide 8 Well Glass Bottom, ibidi) and transfected after 24 h with pc3.1-VHH_enhancer_-HA and pc3.1-mlCNBD-FRET [18] as described above. The medium was replaced with starvation medium (0.5 % FCS) on the following day to induce ciliogenesis. Confocal FRET imaging was performed at the DZNE Light Microscopy Facility using the Andor Spinning Disk Setup (built on an inverted Eclipse Ti Microscope, Nikon) at 37° C. For FRET imaging, the 445 nm laser (18% intensity, 445-, 514-, 640-triple dichroic mirror in the Yokogawa CSU-X1 unit and 5000 rpm disk speed) was used as excitation source, combined with a dual-cam CFP/YFP filter cube (509 nm dichroic mirror with 475/25 nm and 550/49 nm emission filters) to simultaneously measure cerulean and citrine emission with the two EM-CCD cameras (100 ms exposure time, 300 EM gain, 5.36 frames per second frame rate, 10.0 MHz horizontal readout, 1.7 μs vertical readout time, 5x pre Amp gain, −70° C camera temperature). The imaging procedure was as follows: Cells were washed once with ES buffer and measurements were performed in ES buffer. Cilia were imaged with a 100x/1.45 oil objective with 1 μm step size in 10 s intervals. After a stable baseline was obtained, cells were stimulated by drug addition. Cilia-specific fluorescence values were obtained by analyzing the recordings using CiliaQ as described above in “Confocal microscopy and image analysis”. The FRET signal was calculated as a ratio of cerulean/citrine, normalized to the mean baseline value before stimulus addition, and plotted as a change over time.

### ELISA-based cAMP measurements

Total cAMP levels were determined using a CatchPoint^™^ assay (Molecular Devices) according to manufacturer’s instructions. A detailed description can be found in the Supplementary Information.

### Zebrafish as an experimental model

The animal facilities and maintenance of the zebrafish, *Danio rerio*, were approved by the Norwegian Food Safety Authority (NFSA). Detailed information can be found in the SI Appendix. For experiments the following zebrafish lines were used: nacre (*mitfa*-/-) [40], *b-actin:arl13b-gfp* [42], and *Ubi:zebrabow* [44] transgenic animals, which express RFP ubiquitously in absence of Cre recombinase.

### mRNA synthesis, injection, immunostaining, and imaging

The mRNA synthesis, injection, immunostaining, and imaging was performed according to standard protocols. Details are described in the Supplementary Information.

### Data availability statement

The datasets generated during and/or analyzed during the current study are available through the following doi 10.6084/m9.figshare.c.4792248.

### Code availability statement

The analysis workflow to study cilia length and fluorescence signal with its custom-written ImageJ plug-ins (“CiliaQ”) is available through the following link https://github.com/hansenjn/CiliaQ.

