## Supplementary Information for "Nanobody-directed targeting of optogenetic tools to study signaling in the primary cilium"

**Material and methods**

**Plasmids**

The cDNA sequence encoding the anti-mCherry nanobodies was fused to the 3' end of the sequence encoding amino acid 1-201 of mNphp3 (NCBI NM_028721, release 16/09/2018), and to eGFP or an hemagglutinin HA-tag at the 3' end. Similarly, the cDNA encoding the anti-eGFP nanobody was fused to the 5' end of the sequence encoding amino acid 1-201 of mNphp3 and at the 3' end to an HA-tag. The respective constructs were cloned into the pcDNA3.1(+) vector (Thermo Fisher Scientific) for expression in mammalian cells. All primer sequences used for cloning and the corresponding plasmids are summarized in Supplementary Table 1.

**Cell lines and tissue culture**

HEK293 cells were maintained in Dulbecco's modified Eagle's medium (DMEM) (Gibco), supplemented with 1x GlutaMax (Gibco) and 10 % Fetal Calf Serum (FCS) (Biochrome) at 37°C and 5 % $CO_2$ atmosphere. mIMCD-3 were maintained in DMEM/F12 (1:1) medium, supplemented with GlutaMax and 10 % FCS at 37°C and 5 % $CO_2$. Additionally, individual media contained the following: HEK-TM cells: 0.1 mg/ml hygromycin (Thermo Fisher Scientific), HEK-TM bPAC-mCherry: 0.1 mg/ml hygromycin (Thermo Fisher Scientific) and 0.8 mg/ml G418 (Thermo Fisher Scientific), HEK-TM LAPD-mCherry: 50 µg/ml hygromycin, 5 µg/ml blasticidin (Thermo Fisher Scientific), HEK-TM mNphp3(201)-LAPD-mCherry: 0.1 mg/ml hygromycin (Thermo Fisher Scientific) and 0.8 mg/ml G418 (Thermo Fisher Scientific), HEK-mlCNBD-FRET cells: 0.8 mg/ml G418 (Thermo Fisher Scientific), mIMCD-3 bPAC-mCherry: 0.8 mg/ml G418 (Thermo Fisher Scientific), mIMCD-3 LAPD-mCherry: 5 µg/ml blasticidin (Thermo Fisher Scientific). During the experiments, cells were kept without antibiotics.

**Transfection**

For transfection with Lipofectamine 2000 Reagent, the transfection medium was replaced after 4-5 h with full medium. For PEI transfection (4-well dish), 0.5 µg plasmid DNA per well was mixed with 1 µg PEI in 50 µl OptiMEM (Gibco), incubated at room temperature for 10 min, and added to 200 µl full medium on the cells. For PEI transfection (96-well plate), 0.1 µg DNA was mixed with 0.2 µg PEI in 10 µl OptiMEM, incubated at room temperature for 10 min, and added to the cells in 50 µl full medium containing 2% FCS.

**Immunocytochemistry**

Cells were seeded on poly-L-lysine (PLL, 0.1 mg/ml, Sigma Aldrich)-coated 13 mm glass coverslips (VWR) in a 4-well dish (VWR) and transfected on the next day as described above.

For mIMCD-3 cells, the medium was replaced with starvation medium (0.5 % FCS) on the next day to induce ciliogenesis. Cells were fixed 24 h after inducing ciliogenesis (mIMCD-3) or 24-48 h after transfection (HEK293) with 4 % paraformaldehyde (Alfa Aesar, Thermo Fisher Scientific) for 10 min at room temperature. After washing with PBS, cells were blocked with CT (0.5% Triton X-100 (Sigma Aldrich) and 5% ChemiBLOCKER (Merck Millipore) in 0.1 M NaP, pH 7.0) for 30 minutes at room temperature. Primary and secondary antibodies were diluted in CT and incubated for 45 and 60 min at room temperature, respectively. Coverslips were mounted with one drop of Aqua-Poly/Mount (Tebu-Bio).The following antibodies were used: mouse anti-acetylated-Tubulin (1:600, Sigma Aldrich, T6793), rabbit anti-GFP (1:500, Abcam, ab6556, IgG), mouse anti-Arl13B (Abcam, ab136648, 1:500), rabbit anti-Arl13B (1:500, Proteintech, 17711-1-AP), donkey anti-mouse-Cy5 (1:500, Dianova, 715-175-151), goat anti-rabbit-Alexa488 (1:500, Life Technologies, A11034). As a DNA counterstain, DAPI was used (4',6-Diamidino-2-Phenylindole, Dihydrochloride, 1:10 000, Invitrogen) and cells were mounted in Aqua-Poly/Mount (Tebu-Bio).

**Confocal microscopy and image analysis**

For quantifying cilia length and fluorescence signals, z-stacks were recorded from at least two (for stable cell lines) or three (for transiently transfected cells) random positions per experiment and analyzed using custom-written ImageJ plug-ins. Channels were split and the channel representing the Arl13B-staining was segmented by applying an intensity threshold calculated in a maximum projection of the channel ("RenyiEntropy"-algorithm, implemented in ImageJ). The segmented Arl13B channel served as a mask for cilia in the other channels, which were then subjected to a custom-written ImageJ plug-in called "CiliaQ". We developed CiliaQ to fully-automatically quantify the cilia length and ciliary intensity levels in the different channels. CiliaQ detects individual 3D objects in the segmented channel and filters out 3D objects below a pre-defined size threshold (10 voxel) to exclude noise. Each remaining 3D object is considered as a cilium. For each cilium, CiliaQ determines the average intensity of all pixels belonging to the 3D region in each channel. The length of the cilium is determined as the length of the ciliary 3D skeleton obtained by skeletonizing [1] the three-fold-upscaled and blurred (Gaussian blur, sigma = 3 corresponding to 0.21 µm) image of the corresponding ciliary 3D region. All results were scrutinized by a trained observer.

For quantifying centrosomal eGFP fluorescence, z-stacks at nine random positions on the cover slip were recorded and analyzed as described for cilia with the exception that the pericentrin-depicting channel instead of an Arl13B-depicting channel served as a template to reconstruct 3D objects.

**$Ca^{2+}$ imaging**

For probing LAPD activity, HEK-TM or HEK-TM-LAPD cells were seeded on a PLL (0.1 mg/ml, Sigma Aldrich)-coated 96-well plate (F-Bottom, CELLSTAR, Greiner) at 3 x $10^4$ cells per well and incubated over night at 37°C and 5 % $CO_2$ in darkness. For probing bPAC activity, HEK-TM or HEK-TM-bPAC cells were seeded on a PLL (0.1 mg/ml, Sigma Aldrich)-coated 96-well plate (F-Bottom, CELLSTAR, Greiner) at 4 x $10^4$ cells per well, HEK-TM cells were PEI-transfected with pc3-mNphp3(201)-bPAC-mCherry or pcDNA3.1zeo_mCherry on the next day, and incubated over night at 37°C and 5 % $CO_2$ in darkness. For probing LAPD or bPAC activity during nanobody binding, HEK-TM, HEK-TM-LAPD, or HEK-TM-bPAC cells were as described for LAPD activity measurements, transfected on the next day with pEGFP-N1_sstr3, pcDNA3.1-mNphp3(201)-$VHH_{LaM-2}$-eGFP, pcDNA3.1-mNphp3(201)-$VHH_{LaM-2}$-HA, pcDNA3.1-mNphp3(201)-$VHH_{LaM-4}$-eGFP, or pcDNA3.1-mNphp3(201)-$VHH_{LaM-4}$-HA using PEI transfection, and incubated over night at 37°C and 5 % $CO_2$ in darkness. All following steps were conducted under dim green light (LAPD) or dim red light (bPAC). Medium was removed, and cells were washed with 50 µl ES (extracellular solution) buffer (120 mM NaCl, 5 mM KCl, 2 mM $CaCl_2$, 2 mM $MgCl_2$, 10 mM glucose, 10 mM HEPES pH 7.4). Cells were loaded with 2 µM FluoForte (bPAC, bPAC+nanobody, LAPD+nanobody) or 2 µM Fluo4-AM (LAPD) (stocks in DMSO/ Pluronic F-127 (both Sigma-Aldrich)) and 3 mM probenecid (Invitrogen) in 50 µl ES for 30 min at 37°C. Afterwards, the buffer was replaced with 90 µl ES containing 3 mM probenecid, and cells were incubated for 30 min at 29°C in a fluorescence plate-reader (FLUOstar omega, BMG Labtech). Fluorescence was measured at 29°C with an Ex544 excitation and a 570 ± 10 nm emission filter (FluoForte) or with a 485 ± 6 nm excitation and a 530 ± 15 nm emission filter (Fluo4) (all filters BMG Labtech). During bPAC activity measurements, cells were stimulated with a 488-nm-light pulse (1 W/cm²) for 1 s at 3 min, for 10 s at 21 min, and for 60 s at 45 min. For bPAC+nanobody activity measurements, cells were incubated with 25 µM of IBMX (stock: 250 mM in DMSO, AppliChem) from 5 min before light activation on; light activation was induced 2 min after starting the recording using a 488-nm-light pulse (1 s, 162 µW/cm²). For LAPD and LAPD+nanobody activity measurements, cells were stimulated after 2 min with 100 µM NKH477 (Sigma-Aldrich) in ES buffer. During the measurement, the plate was illuminated with an 850-nm LED (0.5 $\mu$W/cm$^2$) inside the reader and then switched to a 690-nm LED (0.5 $\mu$W/cm$^2$) to activate LAPD. At the end of all experiments, ionomycin was added (final concentration: 2 µM, stock: 1 mM in DMSO, Tocris), and fluorescence was recorded until saturation of the signal amplitude. After the end of recording, cell integrity and transfection rate was scrutinized by confocal microscopy of the recorded wells.

**mlCNBD-FRET imaging**

HEK293 or HEK-mlCNBD-FRET cells were seeded and transfected with pcA-Cerulean, pcA-Citrine, pcA-Cerulean + pc3.1-$VHH_{enhancer}$-mCherry, or pcA-Citrine + pc3.1-$VHH_{enhancer}$-

mCherry (HEK293), and pc3.1-VHH$_{enhancer}$-mCherry, pcDNA3.1zeo_mCherry (HEK-mlCNBD-FRET) as described for LAPD activity measurements. Fluorescence imaging of live cells was performed using the CellR Imaging System (Olympus), consisting of an inverse, fully motorized wide-field microscope (IX81) with a monochromatic CCD camera (XM10), a reflector turret, and an illumination system with an excitation-filter wheel (MT20, 150 W Xenon arc burner). Measurements were performed with a 20x/0.75 objective (UPlanSApo, Olympus) at room temperature under atmospheric conditions. Before the measurement, cells were washed once with ES (extracellular solution) buffer and measurements were performed in ES buffer. The experimental recordings were as follows: Before and after each time-resolved measurements, the mCherry fluorescence (12 % light intensity, 200 ms exposure time, 575/25 excitation filter, mCherry-B-0MF Semrock dichroic mirror, 630/20 emission filter) was measured. Time-resolved measurements captured the cerulean (12 % light intensity, 100 ms exposure time, 430/25 excitation filter, M2CFPYFP Olympus dichroic mirror, 480/40 emission filter), the citrine fluorescence (12 % light intensity, 100 ms exposure time, 500/20 excitation filter, M2CFPYFP dichroic mirror, 535-30 emission filter), and the FRET signal (12 % light intensity, 100 ms exposure time, 430/25 excitation filter, M2CFPYFP dichroic mirror, 535-30 emission filter) every 5 s. After 120 s, cells were stimulated with 20 μM isoprenaline hydrochloride (isoproterenol, Sigma Aldrich) or ES buffer as a control. Data was analyzed using Fiji/ImageJ (ImageJ Version 1.52i) [2] by selecting mCherry positive cells with freehand ROIs and determining the mean fluorescence intensity for each ROI in each channel. Values were background subtracted and the FRET signal was corrected for bleed-through and cross-excitation with the following formula: $FRET_{corrected} = FRET – \alpha * cerulean – \beta * citrine$ (with α being the determined bleed-through constant and β being the determined cross-excitation constant for the chosen experimental set-up; α and β values were calculated from single cerulean or citrine transfected cells of three independent experiments using the innate ImageJ tool "Coloc 2", and are 0.75 and 0.02, respectively). Data were plotted as a change of cerulean/$FRET_{corrected}$ over time. Data were acquired from n = 3 independent experiments.

**cADDis imaging**

Mouse IMCD-3 cells were seeded as described above. After 24 h, cells were transduced with the ratiometric cilia-targeted cADDis cAMP assay kit (5-HT$_6$-mCherry-cADDis, Montana Molecular). In detail, 25 μl of the BacMAM stock was mixed with 3 μl sodium butyrate (Sigma Aldrich), and 22 μl Opti-MEM™ (ThermoFischer Scientific). The growth medium on the cells was exchanged with 250 μl Opti-MEM™, and the 50 μl mixture containing BacMAM was added dropwise to the well. Cells were incubated for 24 h at 37° C, 5% $CO_2$ and subsequently measured at the DZNE Light Microscopy Facility using the Andor Spinning Disk Setup. For ratiometric cADDis imaging, cpGFP and mCherry were excited with the 448 nm (10%) and 561 nm (10%) lasers, respectively, in combination with a 405, 448-, 561-, 640-quad dichroic mirror in the Yokogawa CSU-X1 unit and 5000 rpm disk speed. Images were acquired on the two EM-CCD cameras simultaneously (100 ms 100 ms exposure time, 300 EM gain, 5.36 frames per second frame rate, 10.0 MHz horizontal readout, 1.7 µs vertical readout time, 5x pre Amp gain, -70° C camera temperature) with a GFP/RFP emission filter cube (580 nm LP dichroic and 617/73 nm and 525/50 nm emission filters). The experimental procedure during imaging and data analysis was performed as described in the Materials and Methods section "Cilia Imaging". Data analysis was as described above form mlCNBD-FRET sensor imaging, but without correction for bleed-through and cross-excitation. Accordingly, the cADDis signal was plotted as a ratio of mCherry/cpGFP, normalized to the mean baseline value before stimulus addition, and plotted as a change over time.

For imaging in combination with bPAC, mIMCD3 cells were transfected with mNphp3(201)-$VHH_{Lam2}$-HA and bPAC-mCherry or mCherry as described above. After 24 h, cells were transduced with the cilia-targeted cADDis cAMP assay kit (5-$HT_6$-cADDis, Montana Molecular) as described above. Six hours after transduction, the medium was replaced with 250 µl starvation medium per well containing 2 µM sodium butyrate and further incubated at 37° C overnight. 24 h post transduction, cells were measured at the Microscopy Core Facility of the Medical Faculty at the University of Bonn using the Visitron VisiScope Spinning Disk Setup (Build on a Zeiss Axio Observer, Zeiss) at 37° C. For non-ratiometric cADDis imaging in combination with bPAC-mCherry, cpGFP and mCherry were excited with the 448 nm (8%) and 561 nm (10%) laser, respectively, in combination with the 405-488-560bs dichroic mirror in the Yokogawa CSU-W1 unit and the 50 µm pinhole disk at 4000 rpm disk speed. A 60x C-Apochromat water objective (NA = 1.2) was used. Images were acquired on two pco.edge sCMOS cameras simultaneously (200 ms exposure time, 2x binning) with a GFP/RFP emission filter cube. The experimental procedure during imaging and image analysis was the same as described above.

**Zebrafish as an experimental model**

The animal facilities and maintenance of the zebrafish, *Danio rerio*, were approved by the Norwegian Food Safety Authority. Fishes were kept in 3.5 l tanks in a Techniplast Zebtech Multilinking system at 28 °C, pH 7 and 700 mSiemens, at a 14:10 h light/dark cycle. Fish were fed dry food (ZEBRAFEED; SPAROS I&D Nutrition in Aquaculture) two times/day and *Artemia nauplii* once a day (Grade0, platinum Label, Argent Laboratories, Redmond, USA). Embryos were maintained in egg water (1.2 g marine salt and 0.1% methylene blue in 20 l RO water) from fertilization to imaging. All procedures were performed on zebrafish embryos in accordance with the directive 2010/63/EU of the European Parliament and the Council of the European Union and the Norwegian Food Safety Authorities.

**mRNA synthesis, injection, immunostaining, and imaging**

In order to generate capped mRNA of the cherry-nanobody, 5 μg of the plasmid pc3.1-mNphp3 (201)-$VHH_{LaM-2}$-eGFP was first linearized using FastDigest BpiI for 30min at 37 °C (Thermo Fisher Scientific, Cat# FD1014). Upon verification of the linearization of the plasmid by gel electrophoresis, the digested plasmid was purified using the QIAquick PCR purification kit (Qiagen, cat #28104) and its DNA concentration measured using a Nanodrop spectrophotometer (Thermo Fisher Scientific). mRNA was *in vitro* transcribed from 500 ng of linearized plasmid using the mMessage mMachine T7 kit according to the supplier's instructions (Thermo Fisher Scientific , Cat # AM1344). Following 3h incubation at 37 °C, the mRNA was precipitated upon addition of 40 μl RNase-free $H_2O$ and 30 μl LiCl precipitation solution provided in the mMessage mMachine T7 kit, incubation overnight at -20 °C, and centrifugation for 30 min at 13.000 rpm at 4 °C. The precipitated mRNA was further washed with 70% Ethanol in RNase-free $H_2O$, air dried, and resuspended in RNase-free $H_2O$. The integrity of the mRNA was confirmed by gel electrophoresis and its RNA concentration measured with a nanodrop spectrophotometer. The mRNA was aliquoted and kept at -80 °C until injection.

200-300 pg of capped cherry-nanobody mRNA diluted in 0.2M KCl and 0.5% phenol red was injected in the yolk of one-cell stage embryos using a pressure microinjector (Eppendorf Femtojet 4i). Injection needles were pulled with a Sutter Instrument Co. Model P-2000, from thin-walled glass capillaries (1.00 mm; VWR), using the following settings: heat = 450, filament = 4, velocity = 50, delay = 225, pull = 100. The needle tip was cut open with forceps. The pressure and time used for the injection were calibrated for each needle using a 0.01 mm calibration slide for microscopy and a drop of mineral oil to perform injections of 1-2 nl.

Dechorionated and euthanized embryos (collected between 24 and 26 hpf) were fixed in 4 % paraformaldehyde solution and 1 % DMSO for 2 h at room temperature. Embryos were washed 3 times 5 min with 0.3 % Triton-X-100 in PBS (PBSTx), permeabilized with 100% acetone for 10 min at -20 °C, washed three times 10 min with 0.3 % PBSTx and blocked in 0.1 % BSA/0.3 % PBSTx for 2 h. Embryos were incubated with an anti-acetylated tubulin antibody (6-11B-1, 1:1000, Sigma Aldrich) overnight at 4 °C, subsequently washed 3 times 1 h with 0.3 % PBSTx, and incubated with the secondary antibody (Alexa-labelled GAM555 plus, Thermo Fisher Scientific, 1:1,000) and an Alexa Fluor 488-coupled anti-GFP antibody (A-21311, 1:1000, Thermo Fisher Scientific ) overnight at 4 °C. Next, the samples were incubated for 2 h with 0.1 % DAPI in 0.3 % PBSTx (Life Technology), washed three times 1h with 0.3 % PBSTx, and transferred to a series of increasing glycerol concentrations (25 % and 50 %).

Stained larvae were stored in 50 % glycerol at 4 °C and imaged using a Zeiss Examiner Z1 confocal microscope with a 20x plan NA 0.8 objective.

Embryos at 22-26 hpf were manually dechorionated, anaesthetized in 0.01% pH 7.4 buffered MS-222, and mounted in 2% low-melting point agarose (dissolved in artificial fish water (1.2 g marine salt in 20 l RO water) in a Fluorodish (World Precision Instruments). Images were acquired using a Zeiss Examiner Z1 confocal microscope with a 20x water immersion NA 1.0 objective. Acquired images were processed with Fiji/ImageJ (70) or Zen (Zeiss).

**ELISA-based cAMP measurements**

mIMCD-3, mIMCD-3 bPAC-mCherry, or mIMCD-3 LAPD-mCherry cells were seeded on a PLL (0.1 mg/ml, Sigma Aldrich)-coated 96-well plate (F-Bottom, CELLSTAR, Greiner) at $1.8 \times 10^4$ cells per well and incubated over night at 37°C and 5 % $CO_2$ in darkness. During all further experimental procedures, cells or cell lysates were kept in the dark and handled only under dim red (bPAC) or green (LAPD) light, preventing bPAC- or LAPD-activation, respectively. After 48 h, the medium was changed to starvation medium (0.5 % FCS) to induce ciliogenesis. Another 24 h later, the medium was replaced with ES and cells were either subjected to a 2 min light pulse (465 nm, 38.8 µW/cm²; LAPD: 630 nm, 42.3 µW/cm²) or kept in the dark. Directly after light stimulation, cells were lysed and cAMP amounts per well were determined using a CatchPoint™ assay (Molecular Devices), according to manufacturer's instructions. The protein concentration per well was determined with a Pierce™ BCA Protein Assay Kit (Thermo Fisher Scientific), according to manufacturer's instructions.

**Software**

Data analysis and statistical analysis was performed in Excel (Microsoft Office Professional Plus 2013, Microsoft) and GraphPad Prism (Version 8.1.2, GraphPad Software, Inc.). All image processing and analysis was performed in ImageJ (Version v1.52i, U.S. National Institutes of Health, Bethesda, Maryland, USA). Plots and Figures were generated using GraphPad Prism (Version 8.1.2, GraphPad Software, Inc.) and Adobe Illustrator CS5 (Version v15.0.0, Adobe Systems, Inc.). ImageJ plugins were developed in Java, with the aid of Eclipse Mars.2 (Release 4.5.2, IDE for Java Developers, Eclipse Foundation, Inc., Ottawa, Ontario, Canada).

### Supplementary Figures

#### Supplementary Figure 1

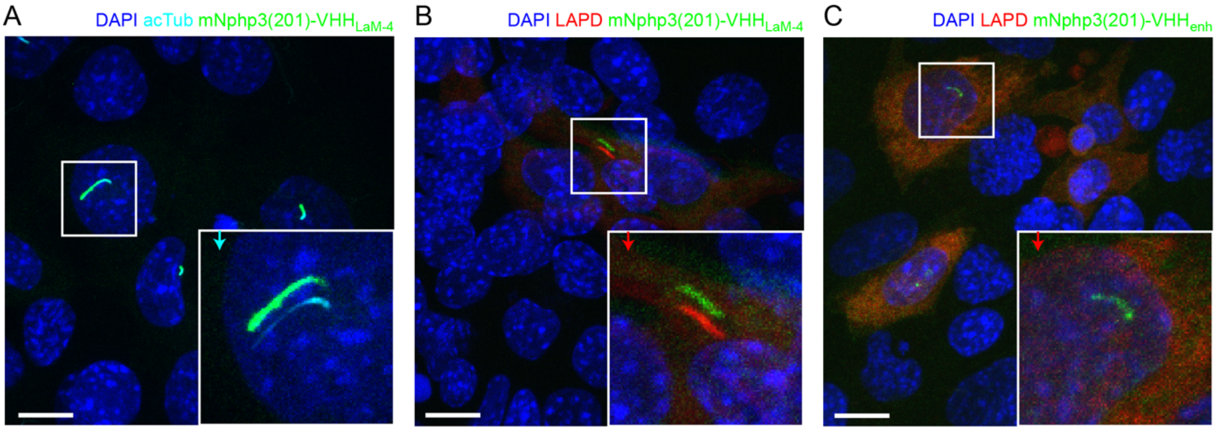


**Supplementary Figure 1: Nanobody-based ciliary targeting. A.** Localization of the anti-mCherry nanobody to primary cilia. mIMCD-3 cells were transfected with mNphp3(201)-$VHH_{LaM-4}$ and stained with an acetylated tubulin antibody (cyan, ciliary marker) and DAPI (blue). **B.** Localization of the anti-mCherry nanobody and LAPD-mCherry to primary cilia. mIMCD-3 cells were co-transfected with mNphp3(201)-$VHH_{LaM-4}$-eGFP (green) and LAPD-mCherry (red) and labeled with DAPI (blue). **C.** Localization of the anti-GFP nanobody mNphp3(201)-$VHH_{enhancer}$-HA to primary cilia, while LAPD-mCherry resides in the soma. mIMCD-3 cells were co-transfected with mNphp3(201)-$VHH_{enhancer}$-HA (green) and LAPD-mCherry (red) and stained with an anti-HA antibody (green) and DAPI (blue). Boxes indicate the position of the magnified view shown at the bottom right. Arrows in different colors indicate the direction and the length of the shift of the respective fluorescence channel. Scale bars: 10 µm.

### Supplementary Figure 2

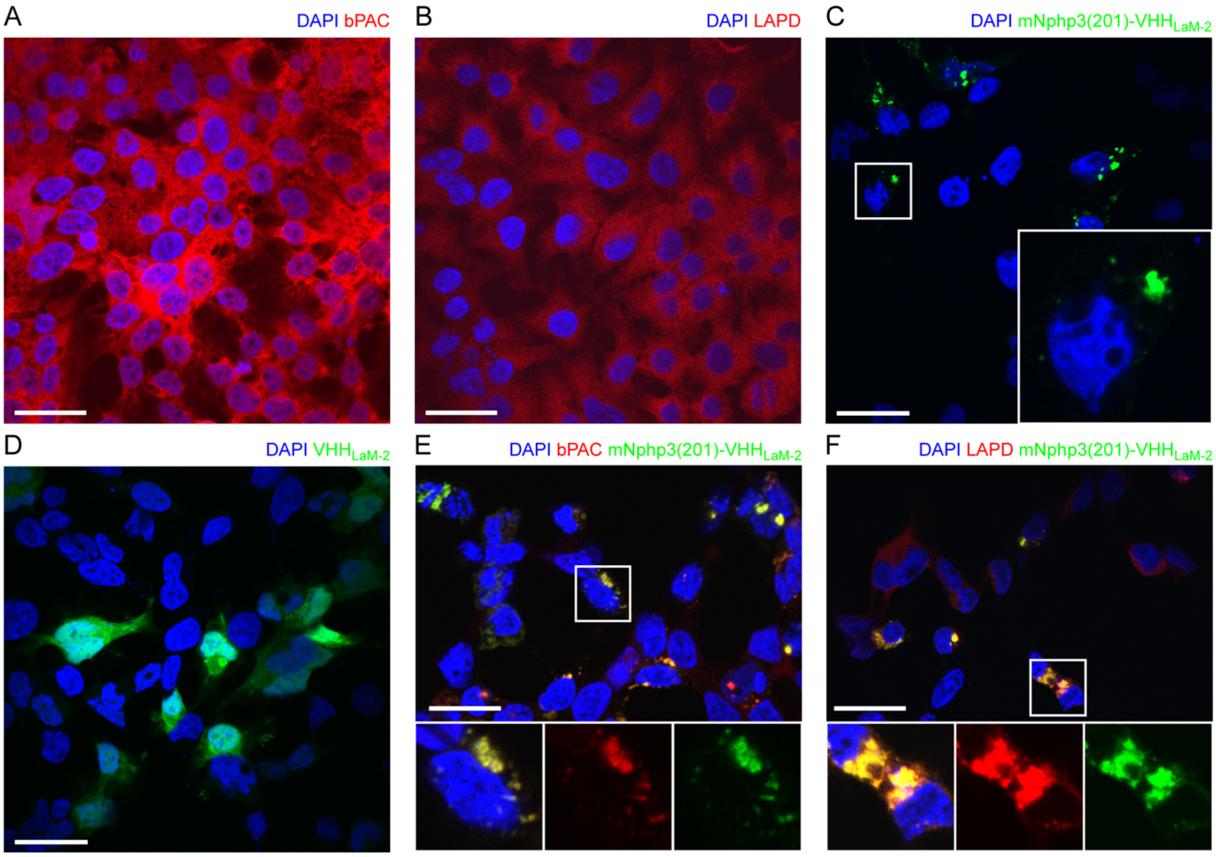


**Supplementary Figure 2: Subcelluar localization of nanobody-targeted optogenetic tools. A.** HEK-TM cells expressing bPAC. HEK-TM cells stably expressing bPAC-mCherry (red) were fixed and labeled with DAPI (blue). Scale bar: 20 μm. **B.** See A. for LAPD-mCherry. **C.** HEK-TM cells expressing the cilia-targeted anti-mCherry mNphp3(201)-$VHH_{LaM-2}$-eGFP (green) nanobody. The box indicates the position of the magnified view shown at the bottom right. **D.** HEK-TM cells expressing the cilia-targeted anti-mCherry $VHH_{LaM-2}$-eGFP (green) nanobody. **E.** Co-expression of the cilia-targeted anti-mCherry nanobody (LaM-2) (green) and bPAC-mCherry (red) in HEK-TM cells. The box indicates the position of the magnified view shown at the bottom: left: all channels as overlay; center: mCherry channel only; right: eGFP channel only. **F.** Co-expression of the cilia-targeted anti-mCherry nanobody (LaM-2) (green) and LAPD-mCherry (red) in HEK-TM cells. The box indicates the position of the magnified view shown at the bottom: left: all channels as overlay; center: mCherry channel only; right: eGFP channel only.

**Supplementary Figure 3**

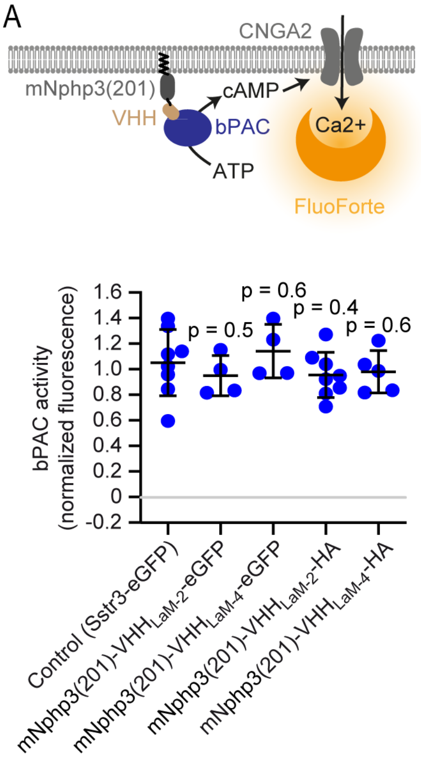


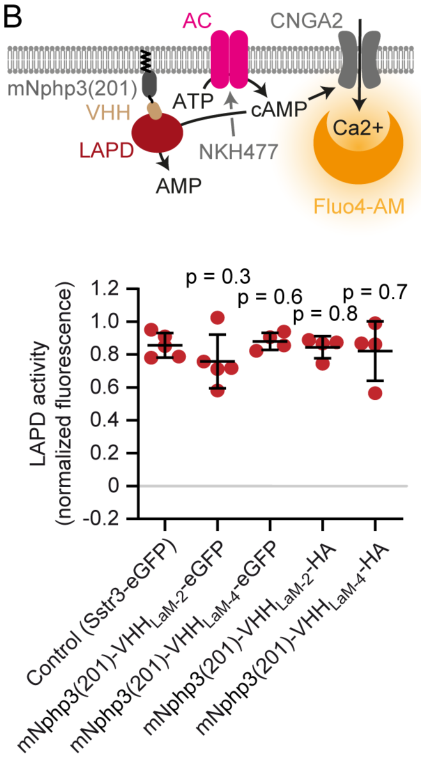


**Supplementary Figure 3: Activity measurements in HEK-TM cells. A.** bPAC activity measurements in HEK-TM cells. HEK293 cells express the CNGA2-TM ion channel, which opens upon cAMP binding and conducts $Ca^{2+}$ (HEK-TM) [3]. bPAC-mCherry was co-expressed with the mNphp3(201)-tagged mCherry nanobody. Light-dependent activation of bPAC increases intracellular cAMP levels, leading to a $Ca^{2+}$ influx, which was quantified using a fluorescent $Ca^{2+}$ dye (GFP-certified FluoForte). bPAC activity was determined in the presence of mNphp3(201)-$VHH_{LaM-2}$ or mNphp3(201)-$VHH_{LaM-4}$ (fused to HA or eGFP). Co-expression with the ciliary protein Sstr3-eGFP was used as a negative control (ciliary localized protein, but not binding to bPAC or LAPD). bPAC activity was determined according to the maximum amplitude of the $Ca^{2+}$ signal after light stimulation (465 nm light pulse, 1 sec, 162 µW/cm²) compared to the ionomycin-evoked $Ca^{2+}$ signal. 5 min before light stimulation, cells were treated with 25 µM of IBMX to inhibit phosphodiesterases and sustain a long-lasting increase in cAMP. NT: non-transfected cells. **B.** LAPD activity measurements in HEK-TM cells. To measure LAPD activity, HEK-TM cells were pre-stimulated with 100 µM NKH477 to activate transmembrane adenylate cyclases (AC), thus increasing cAMP levels. $Ca^{2+}$ influx was detected by a $Ca^{2+}$ dye (Fluo4-AM). LAPD activity was determined in the presence of mNphp3(201)-$VHH_{LaM-2}$ or mNphp3(201)-$VHH_{LaM-4}$ (fused to HA or eGFP). Co-expression of the ciliary protein Sstr3-eGFP was used as a negative control (ciliary localized protein, but not binding to bPAC or LAPD).  Fluo4-AM-loaded HEK-TM cells were incubated with 100 µM NKH477 during continuous 850 nm light illumination (0.5 µW/cm²). When reaching a steady-state, light was switched to 690 nm (0.5 µW/cm²) to stimulate LAPD activity. LAPD activity was determined as the maximal decrease compared to the maximal $Ca^{2+}$ signal amplitude after NKH477 addition. Data are shown as individual data points (each data point represents and independent experiment and corresponds to the average of a duplicate or triplicate measurement) and mean ± S.D., p-values calculated using unpaired, two-sided Student's t-test compared to Sstr3-eGFP are indicated. All HEK-293 cells were non-ciliated.

### Supplementary Figure 4

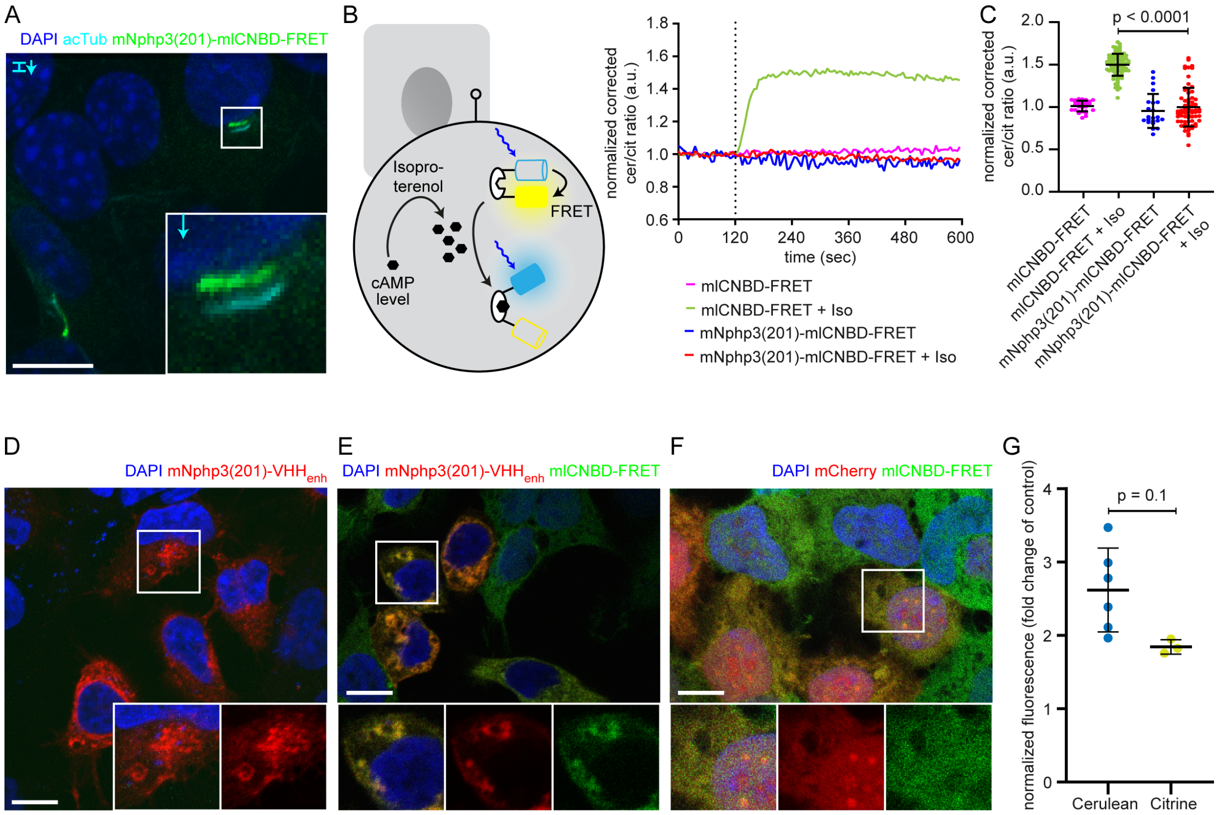


**Supplementary Figure 4: Characterization of the ciliary-targeted cAMP mlCNBD-FRET biosensor. A.** Localization of mNphp3(201)-mlCNBD-FRET to primary cilia in mIMCD-3 cells. **B.** FRET imaging in non-ciliated HEK293 cells expressing mlCNBD-FRET or mNphp3(201)-mlCNBD-FRET under control conditions or after stimulation with 20 μM isoproterenol (addition depicted with dotted line). Data are shown as mean (n = 3 independent experiments, 30-90 cells per experiment). Schematic overview of mlCNBD-FRET imaging is shown on the left. **C.** Comparison of the maximal FRET change for data shown in B. Data are presented as individual data points and mean ± S.D.; p-value calculated using a two-tailed Mann-Whitney test is indicated. **D.** Localization of mNphp3(201)-$VHH_{enhancer}$-mCherry in HEK293 cells. The box indicates the position of the magnified view shown at the bottom for the individual channels. **E./F.** Localization of mlCNBD-FRET in HEK293 cells in the presence (E) or absence (F) of mNphp3(201)-$VHH_{enhancer}$-mCherry. In F, mCherry only was used as a control. Scale bar: 10 μm. The box in indicates the position of the magnified view shown at the bottom. left: overlay; middle and right: individual channels. **G.** HEK293 cells were transfected with cerulean or citrine in the presence of mCherry (control) or the eGFP nanobody mNphp3(201)-$VHH_{enhancer}$-mCherry. Fluorescence intensities of cerulean or citrine were normalized to the mCherry fluorescence of the nanobody or mCherry only in the same cell, and the relative change in fluorescence compared to the control condition (mCherry only) was plotted. Data are shown as mean ± S.D., n = 3-6 with 2-30 cells per experiment; p-values were determined using an unpaired, two-sided Student's t-test.

### Supplementary Figure 5

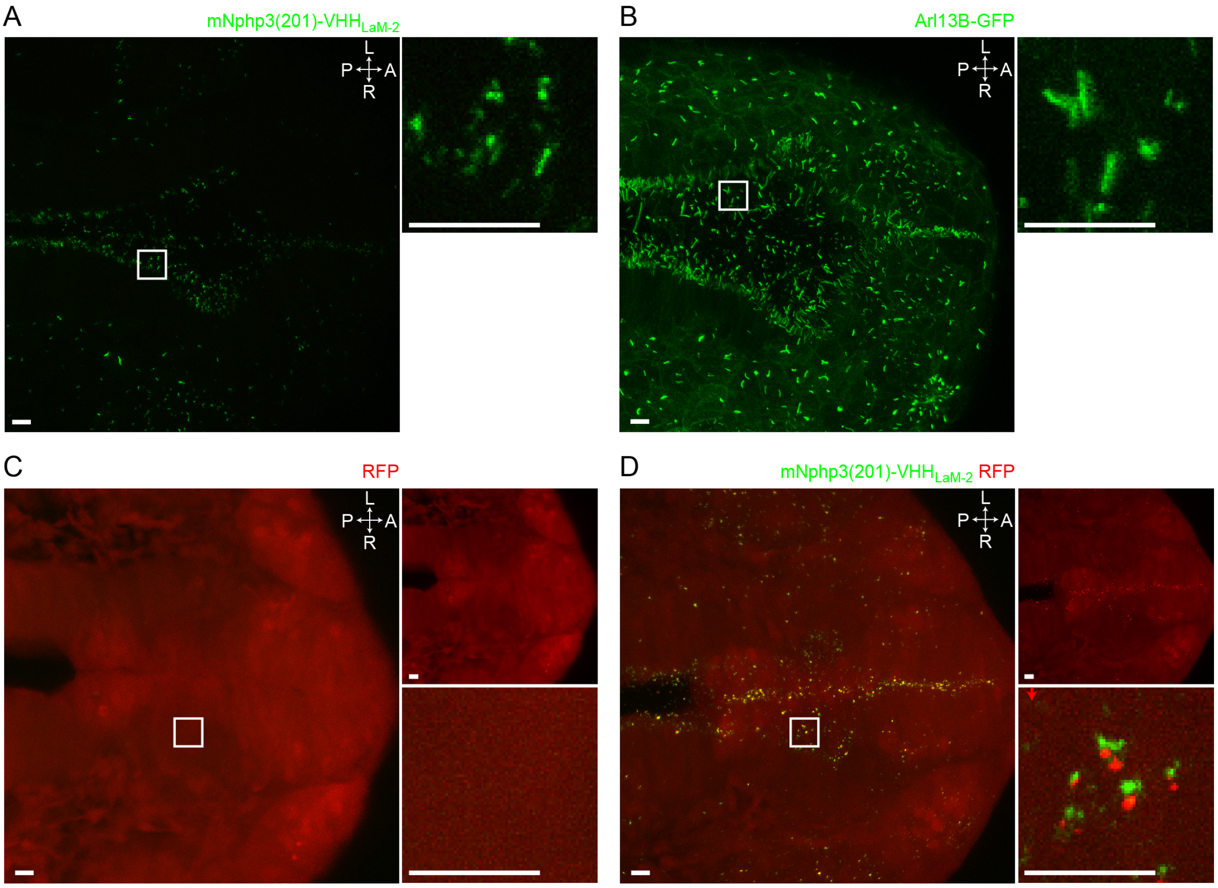


**Supplementary Figure 5: Nanobody-based ciliary protein targeting *in vivo.* A.** Nanobody localization in the neural tube of a zebrafish embryo. The mRNA of the anti-mCherry mNphp3(201)-VHH$_{LaM-2}$-eGFP (green) nanobody was injected into nacre zebrafish embryos. **B.** Ciliary labeling (green) in the neural tube of b-actin:arl13b-gfp transgenic embryos (38). **C.** RFP (red) expression in the neural tube of Ubi:zebrabow (40) transgenic embryos. **D.** RFP (red) expression in the neural tube of Ubi:zebrabow (40) transgenic embryos, injected with mRNA of the anti-mCherry mNphp3(201)-VHH$_{LaM-2}$-eGFP nanobody. Scale bar: 20 μm, magnified view: 10 μm. Boxes indicate the position of the magnified views (bottom right panels). Arrow indicates the direction and the length of the shift of the respective fluorescence channel. The upper right panels show the individual RFP fluorescence, the bottom magnified view the overlay. A: anterior, P: posterior, L: left, R: right, D: dorsal, V: ventral. All images have been aquired from live embryos.

**Supplementary Figure 6**

A

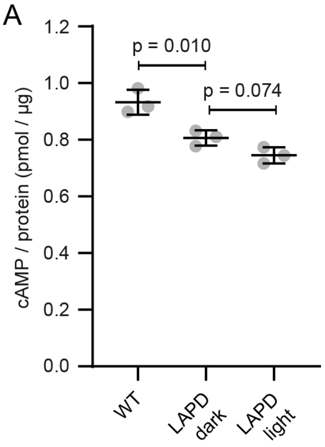


**Supplementary Figure 6: cAMP levels and ciliary length in mIMCD-3 cells expressing LAPD. A.** Determination of total cAMP levels in WT mIMCD-3 cells or mIMCD-3 LAPD-mCherry cells in the dark or after light stimulation (2 min, 630 nm, 42.3 µW/cm$^2$). cAMP levels have been determined using an ELISA and normalized to the protein concentration. Data are shown as mean ± S.D., n = 3, p-values were calculated using a paired two-sided Student's t-test.

**Supplementary Table 1**

**Supplementary Table 1: Plasmids and cloning information.** Plasmids are listed and the ID and sequence of the primers that have been used for cloning are indicated.

| **pcDNA3.1_bPAC-mCherry** | | |
|---|---|---|
| **ID** | **Sequence 5'-3'** | **Primer Info** |
| C0654 | CCGGATCCACCATGATGAAGCGGCTGGTGTAC | 5' primer for PCR bPAC, adds BamHI |
| C0631 | CCCAAGCTTGGCGCGCCGGCAGGCGCCACTTGGC | 3' primer for PCR bPAC, adds AscI |
| C0633 | CCCAAGCTTACTTGTACAGCTCGTCCATG | 3' primer for PCR mCherry, adds HindIII |
| **pc6-LAPD-mCherry** | | |
| **ID** | **Sequence** | **Primer Info** |
| C2041 | GGATCCTAAGCTTCCACCATGAGCAGGGACCCCC | 5' primer for PCR LAPD, adds HindIII/Kozak |
| C1724 | GGTTCTAGATTACTTGTACAGCTCGTCCATGCC | 3' primer for PCR mCherry, adds XbaI |
| **pc3-mNphp3(201)-bPAC-mCherry** | | |
| **ID** | **Sequence 5'-3'** | **Primer Info** |
| C2956 | GGAGGATCCATGATGAAGCGGCTGGTGTAC | 5' primer for PCR of mNPHP3, adds BamHI |
| C2954 | GTAGTCGGGCACGTCGTAGGGGTACTTGTACAGCTCGTCCATGCCG | 3' primer for PCR mCherry, adds HA |
| C0609 | TCTTCTAGATTAGGCGTAGTCGGGCACGTCGTAGGGG | 3' primer for PCR mCherry, adds HA, XbaI |
| **pc3-mNphp3(201)-LAPD-mCherry** | | |
| **ID** | **Sequence 5'-3'** | **Primer Info** |
| C2951 | GGAGGATCCATGAGCAGGGACCCCCTGCC | 5' primer for PCR of mNPHP3, adds BamHI |
| C2952 | CACCATCGTCGCGACCGGTGGGTCCCGGGCCCGCGGTAC | 3' primer for PCR of mNPHP3+linker, adds PVAT linker |
| C2953 | GACCCACCGGTCGCGACGATGGTG | 5' primer for PCR of mNPHP3+linker, adds PVAT linker |
| C0609 | TCTTCTAGATTAGGCGTAGTCGGGCACGTCGTAGGGG | 3' primer for PCR mCherry, adds HA, XbaI |

| | | |
|---|---|---|
| **pcDNA3.1zeo_mCherry** | | |
| **ID** | **Sequence 5’-3’** | **Primer Info** |
| 5007 | GAAGGATCCACCATGGTGAGCAAGG GCGAGG | 5’ primer for PCR mCherry, adds BamHI |
| 5008 | CTCTCTAGATTACTTGTACAGCTCGT CCATG | 3’ primer for PCR mCherry, adds XbaI |
| **pc3-mNphp3(201)-mCherry** | | |
| **ID** | **Sequence 5’-3’** | **Primer Info** |
| C3037 | CGACGATGACGATAAGGATCCATGG TGAGCAAGGGCGAGG | 5’ primer for PCR of mNPHP3, adds BamHI |
| C2637 | ATATCTAGATTACTTGTACAGCTCGT CCATGCC | 3’ primer for mCherry, adds XbaI |
| **pcA-Cerulean** | | |
| **ID** | **Sequence 5’-3’** | **Primer Info** |
| 4037 | CGGGATCCACCATGGTGAGCAAGGG CGAG | 5’ primer for PCR of cerulean/citrine, adds BamHI |
| 4036 | GCTCTAGATTACTTGTACAGCTCGTC CATGCC | 3’ primer for PCR of cerulean/citrine, adds XbaI |
| **pcA-Citrine** | | |
| **ID** | **Sequence 5’-3’** | **Primer Info** |
| 4037 | CGGGATCCACCATGGTGAGCAAGGG CGAG | 5’ primer for PCR of cerulean/citrine, adds BamHI |
| 4036 | GCTCTAGATTACTTGTACAGCTCGTC CATGCC | 3’ primer for PCR of cerulean/citrine, adds XbaI |
| **pcDNA3.1-mNphp3(201)-$\text{VHH}_{\text{LaM-2}}$-eGFP** | | |
| **ID** | **Sequence 5’-3’** | **Primer Info** |
| C4052 | ATTACTCGAG TGAGGAGACG GTGACCTGGG | 3' primer for PCR of VHH, XhoI in frame for fusion to eGFP |
| C4053 | TAATCTCGAG ATGGTGAGCA AGGGCGAGGA | 5' primer for PCR of eGFP, XhoI in frame for fusion to VHH |
| C4054 | ATTCTCTAGA TTACTTGTAC AGCTCGTCCA TGCC | 3' primer for PCR of eGFP, XbaI after Stop, for cloning eGFP to VHH |
| C4095 | TATAGGATCC CAGGTGCAGC TCGTGGAAAG TG | 5' primer for PCR of VHH-LaM-2, adds BamHI for fusion to mNPHP3 and adds Kozak |
| **pcDNA3.1-mNphp3(201)-$\text{VHH}_{\text{LaM-2}}$-HA** | | |

| ID | Sequence 5’-3’ | Primer Info |
|---|---|---|
| C4095 | TATAGGATCC CAGGTGCAGC TCGTGGAAAG TG | 5' primer for PCR of VHH-LaM-2, adds BamHI for fusion to mNPHP3 and adds Kozak |
| C4144 | GGTCTAGATC ACGCATAATC CGGCACATCA TACGGATATG AGGAGACGGT GACCTGG | 3' primer for PCR of VHH with HA tag at 3' end, adds Stop codon and XbaI for ligation into vector |
| **pcDNA3.1-VHH$_{LaM-2}$-eGFP** | | |
| **ID** | **Sequence 5’-3’** | **Primer Info** |
| C4096 | TATAGGATCC ACCATGCAGG TGCAGCTCGT GGAAAGTG | 5' primer for PCR of VHH-Lam2, adds BamHI for cloning into vector, Kozak, ATG |
| C4052 | ATTACTCGAG TGAGGAGACG GTGACCTGGG | 3' primer for PCR of VHH, XhoI in frame for fusion to eGFP |
| C4053 | TAATCTCGAG ATGGTGAGCA AGGGCGAGGA | 5' primer for PCR of eGFP, XhoI in frame for fusion to VHH |
| C4054 | ATTCTCTAGA TTACTTGTAC AGCTCGTCCA TGCC | 3' primer for PCR of eGFP, XbaI after Stop, for cloning eGFP to VHH |
| **pcDNA3.1-mNphp3(201)-VHH$_{LaM-4}$-eGFP** | | |
| **ID** | **Sequence 5’-3’** | **Primer Info** |
| C4051 | TATAGGATCC CAGGTGCAGC TCGTGGAATC TG | 5' primer for PCR of VHH, BamHI in frame for fusion to mNphp3(201) |
| C4052 | ATTACTCGAG TGAGGAGACG GTGACCTGGG | 3' primer for PCR of VHH, XhoI in frame for fusion to eGFP |
| C4053 | TAATCTCGAG ATGGTGAGCA AGGGCGAGGA | 5' primer for PCR of eGFP, XhoI in frame for fusion to VHH |
| C4054 | ATTCTCTAGA TTACTTGTAC AGCTCGTCCA TGCC | 3' primer for PCR of eGFP, XbaI after Stop, for cloning eGFP to VHH |
| **pcDNA3.1-mNphp3(201)-VHH$_{LaM-4}$-HA** | | |
| **ID** | **Sequence 5’-3’** | **Primer Info** |
| C4051 | TATAGGATCC CAGGTGCAGC TCGTGGAATC TG | 5' primer for PCR of VHH, BamHI in frame for fusion to mNphp3(201) |
| C4144 | GGTCTAGATC ACGCATAATC CGGCACATCA TACGGATATG AGGAGACGGT GACCTGG | 3' primer for PCR of VHH with HA tag at 3' end, adds Stop codon and XbaI for ligation into vector |
| **pcDNA3.1-VHH$_{LaM-4}$-eGFP** | | |
| **ID** | **Sequence 5’-3’** | **Primer Info** |
| C4052 | ATTACTCGAG TGAGGAGACG GTGACCTGGG | 3' primer for PCR of VHH, XhoI in frame for fusion to eGFP |

| | | |
|---|---|---|
| C4053 | TAATCTCGAG ATGGTGAGCA AGGGCGAGGA | 5' primer for PCR of eGFP, XhoI in frame for fusion to VHH |
| C4054 | ATTCTCTAGA TTACTTGTAC AGCTCGTCCA TGCC | 3' primer for PCR of eGFP, XbaI after Stop, for cloning eGFP to VHH |
| C4061 | TATAGGATCC ACCATGCAGG TGCAGCTCGT GGAATCTG | 5' primer for PCR of VHH-LaM-4, adds BamHI for cloning into vector, Kozak, and ATG |
| **pcDNA3.1-mNphp3(201)-VHH$_{enhancer}$-HA** | | |
| **ID** | **Sequence 5'-3'** | **Primer Info** |
| C4142 | AACGGATCCC AGGTGCAGCT GCAGGAATC | 5' primer for PCR of VHH-enhancer, adds BamHI for fusion to Nphp3 and deletes ATG |
| C4143 | GGTCTAGATC AGGCATAATC TGGGACATC | 3' primer cloning VHH-enhancer fusion with Nphp3, adds XbaI after HA and stop |
| **pc3.1-mNphp3(201)-VHH$_{enhancer}$-mCherry** | | |
| **ID** | **Sequence 5'-3'** | **Primer Info** |
| C4142 | AACGGATCCC AGGTGCAGCT GCAGGAATC | 5' primer for PCR of VHH-enhancer, adds BamHI for fusion to Nphp3 and deletes ATG |
| C4263 | TAACTCGAGC ATCCCGGGTA CCATGCATCG | 3' primer for PCR of VHH-enhancer, deletes Stop and adds XhoI for fusion to mCherry |
| C4264 | GAACTCGAGG TGAGCAAGGG CGAGGAGGAT | 5' primer for PCR of mCherry, adds XhoI for fusion to VHH-enhancer and deletes ATG |
| C4265 | CGGTCTAGAT CACTTGTACA GCTCGTCCAT GC | 3' primer for PCR of mCherry, adds XbaI after Stop for ligation into vector |
| **pc3.1-VHH$_{enhancer}$-mCherry** | | |
| **ID** | **Sequence 5'-3'** | **Primer Info** |
| C4265 | CGGTCTAGAT CACTTGTACA GCTCGTCCAT GC | 3' primer for PCR of mCherry, adds XbaI after Stop for ligation into vector |
| C4266 | TGAGGATCCA CCATGCAGGT GCAGCTGCAG GAATCGGG | 5' primer for PCR of VHH-enhancer-mCherry, adds BamHI for ligation into vector, Kozak, and ATG |
| **pEGFP-N1-bPAC** | | |
| C4443 | AATCGCTAGCCACCATGAAGCGGCT GGTGTACATC | 5' primer cloning bPAC in pEGFP-N1, adds NheI and Kozak before ATG |
| C4444 | GCCGAATTCTGTTCTTGTCGTTTTCC AGGGTCTGC | 3' primer cloning bPAC in pEGFP-N1, no stop, extra 4 bp, then EcoRI site |
